# The PH domain in the ArfGAP ASAP1 drives catalytic activation through an unprecedented allosteric mechanism

**DOI:** 10.1101/2024.12.20.629688

**Authors:** Olivier Soubias, Samuel L. Foley, Xiaoying Jian, Rebekah A. Jackson, Yue Zhang, Eric M. Rosenberg, Jess Li, Frank Heinrich, Margaret E. Johnson, Alexander J. Sodt, Paul A. Randazzo, R. Andrew Byrd

**Affiliations:** Center for Structural Biology, Center for Cancer Research, National Cancer Institute, National Institutes of Health, Frederick, MD, USA; Department of Biophysics, The Johns Hopkins University, Baltimore, MD, USA; Laboratory of Cellular and Molecular Biology, Center for Cancer Research, National Cancer Institute, National Institutes of Health, Bethesda, MD 20892, USA; Department of Physics Carnegie Mellon University, Pittsburgh, PA, USA; NIST Center for Neutron Research, Gaithersburg, MD, USA; Unit of Membrane Chemical Physics, Eunice Kennedy Shriver National Institute of Child Health and Human Development, National Institutes of Health, Bethesda, MD, USA

## Abstract

ASAP1 is a multidomain Arf GTPase-activating protein (ArfGAP) that catalyzes GTP hydrolysis on the small GTPase Arf1 and is implicated in cancer progression. The PH domain of ASAP1 enhances its activity greater than 7 orders of magnitude but the underlying mechanisms remain poorly understood. Here, we combined Nuclear Magnetic Resonance (NMR), Molecular Dynamic (MD) simulations and mathematical modeling of functional data to build a comprehensive structural-mechanistic model of the complex of Arf1 and the ASAP1 PH domain on a membrane surface. Our results support a new conceptual model in which the PH domain contributes to efficient catalysis not only by membrane recruitment but by acting as a critical component of the catalytic interface, binding Arf·GTP and allosterically driving it towards the catalytic transition state. We discuss the biological implications of these results and how they may apply more broadly to poorly understood membrane-dependent regulatory mechanisms controlling catalysis of the ArfGAP superfamily as well as other peripheral membrane enzymes.

## Introduction

Adenosine diphosphate–ribosylation factors (Arfs) are a family of GTPases that control membrane traffic, cytoskeletal dynamics and lipid signaling. The function of Arfs, which have no detectable intrinsic GTPase activity, is dictated by over 30 Arf guanosine triphosphatase– activating proteins (GAPs) that induce hydrolysis of Arf-bound GTP. The precise roles and the molecular bases for Arf interaction with the GAPs are still being discovered^1,2^.

ASAP1 (ArfGAP with SH3 domain, Ankyrin repeat and PH domain 1) is a 130 kDa multidomain polypeptide composed of BAR (Bin/Amphiphysin/Rvs), PH (Pleckstrin Homology), Arf GAP, Ankyrin Repeat, proline rich, and SH3 (Src Homology 3) domains that controls actin and cell adhesion dynamics and is thought to contribute to invasion and metastasis in cancer^2^. A catalytic arginine in the Arf GAP domain is essential but not sufficient for maximum GTP hydrolysis. The isolated Arf GAP domain has low catalytical efficiency (∼ 1 M^-1^sec^-1^). In contrast, a protein fragment containing both the PH, Arf GAP and Ankyrin Repeat domains acts as a robust GAP, with eight orders of magnitude greater efficiency (∼ 10^8^ M^-1^sec^-1^)^3,4^.

One established role of PH domains is to bind to phosphatidylinositol phosphates (PIPs)and proteins at the membrane^5^, thus restricting PH domain-containing proteins to the volume containing a target peripheral membrane protein and increasing the effective concentration, as well as the frequency of collision^6^. PH domains can also function in other capacities including autoinhibition or positioning other structural elements of a protein to inhibit intramolecular catalytic domains, as described for kinases and guanine nucleotide exchange factors^7–9^, or forming part of the substrate-binding site, as observed in Rho exchange factors such as Dbs and FARP2 (FERM, ARH/ RhoGEF and pleckstrin domain protein 2)^10,11^. While autoinhibition does not appear to be a regulatory mechanism in ASAP1, the molecular mechanism underlying the large PI(4,5)P_2_-dependent increase in enzymatic activity remains unknown. Here, we integrated Nuclear Magnetic Resonance (NMR), Molecular Dynamics (MD) simulations, and mathematical modeling to test hypotheses about the mechanisms by which the PH domain contributes to efficient catalysis. Our analysis indicates that the PH domain enhances catalytic activity by (i) restricting ASAP1 to a membrane surface to accelerate binding to the membrane associated substrate Arf1·GTP, (ii) binding to its substrate Arf1·GTP, providing proximity and orientation to the catalytic residues in the Arf GAP domain, and (iii) remodeling the nucleotide binding site in Arf1·GTP to achieve a transition state intermediate. Thus, the PH domain is an integral component of the catalytic interface. As far as we are aware this is (i) the first report of a PH domain directly contributing catalytic activation of a G protein and (ii) the first quantitative analysis of the relative contribution of membrane recruitment and allosteric activation of a protein by a PH domain. We discuss the ramifications for understanding Arf signaling and for designing small molecule inhibitors of proteins containing PH domains, and whether the paradigm might extend to other PH domain-containing proteins.

## RESULTS

### ASAP1 PH contributes to catalysis beyond membrane recruitment

GAP activity of ASAP1 fragments were previously measured in our laboratories^12–15^. However, since these studies were done under different experimental conditions, we repeated the measurements in the presence of large unilamellar vesicles (LUVs) with or without PI(4,5)P_2_ (5 mol%). The GAP activities of *wt* PZA (**<u>P</u>**H domain, **<u>Z</u>**inc binding and **<u>A</u>**nkyrin repeats, [325-724]ASAP1), ^ΔN14^PZA (where residues 325-338 corresponding to the first 14 residues of the *wt* PH domain are truncated), PHdZA (PH domain of phospholipase Cδ1 in tandem with ASAP1 ZA domain) which all binds to PI(4,5)P_2_ equally well through their PH domain but have almost no affinity for PI(4,5)P_2_ free-membranes^12,15^ and ^ΔPH^ZA domain ([431-724]ASAP1, henceforth called ZA), which lacks the PH domain, were compared (Fig. 1A, B). GAP activities depended on the protein constructs and on the presence of PI(4,5)P_2_ (*wt* PZA^PIP^^2^ > ^ΔN14^PZA^PIP^^2^ > PHdZA^PIP^^2^ > *wt* PZA^PC^ > PHdZA^PC^ ≈ ZA^PIP2^ ^or^ ^PC^) indicating that although PH-dependent membrane recruitment contributes to GAP activity, the cognate PH domain enhances GAP activity by other mechanisms, including binding to Arf (Fig1. C,D and Table SI1).

**Fig. 1.**
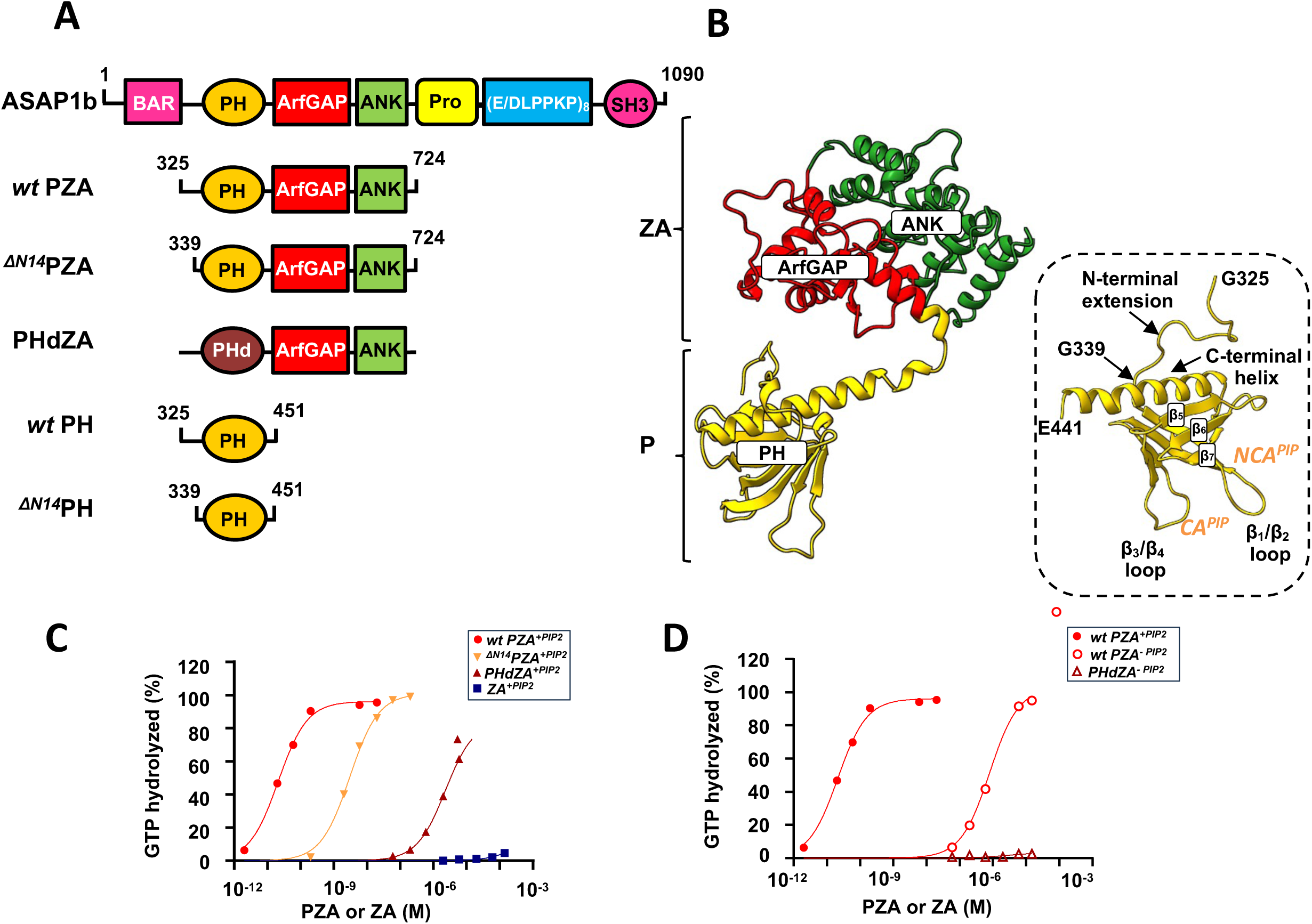
Binding of ASAP1 PH to myrArf1 is key for maximal GTP hydrolysis. A) Left. Schematic of recombinant proteins used in this paper. The domain structure of ASAP1 is shown in the schematic at top. Abbreviations: BAR, Bin/amphiphysin/RVS; PH, pleckstrin homology; Arf GAP, Arf GTPase-activating protein; ANK, ankyrin repeat; Pro-Rich, proline-rich; (E/DLPPKP)8, tandem repeats of E/DLPPKP; SH3, Src homology 3. Recombinant proteins used in the studies are shown below the schematic of full-length ASAP1. The acronyms for the proteins include “P” for the PH domain, “Z” for the Arf GAP domain, which is a zinc-binding motif, “A” for the ankyrin repeat and “PHd” for the PH domain of phospholipase Cδ1. PHdZA is a chimeric protein consisting of residues 1 to 134 of PLCδ1 and residues 441 to 724 of ASAP1 **B)** Ribbon representation of the AlphaFold structure of *wt* PZA. The PH (in yellow) and ZA (in red/green) domains behaves like “beads-on-a-string”^57^. Inset: Ribbon representation of the structure of ASAP1–PH in PDB: 5C79. PH domains are defined by a structural fold of 7 β-strands arranged as a sandwich and capped by a C-terminal amphipathic α-helix with loops of different lengths connecting the β-strands. For visual guidance, β strands 5-7, as well as loops linking the β strands and C-terminal helix are labeled. Residues 442-451 (unstructured) are not shown for clarity. Approximate sites of PI(4,5)P_2_ interaction are labeled CA (for canonical sites) and NCA (for noncanonical site). **C)** Comparison of GAP activity using Arf1 as substrate in the presence of PI(4,5)P_2_ containing membranes. *wt* PZA (red circle), ^ΔN14^PZA (orange ▼), PHdZA (maroon ▲) or ZA (blue ▪) was titrated into a GAP reaction containing 1 μM full-length Arf1 and LUVs at a total phospholipid exposed concentration of 0.5 mM containing 5% mol PI(4,5)P_2_. **D)** Comparison of GAP activity using Arf1 as substrate in the presence of membranes without PI(4,5)P_2_ for *wt* PZA (red circle), ^ΔN14^PZA (orange open triangle) and PHdZA (maroon open triangle).

To separate the effect of membrane recruitment from other activating mechanisms, we measured the rate of GTP hydrolysis “*in trans*” by tryptophan fluorescence and in the presence of nanoscale lipid domains (nanodiscs (NDs))^16,17^, using isolated PH and ZA domains^18–21^. In the presence of ZA alone, GTP hydrolysis was not detected. Adding the PH domain to the reaction mixture with ZA triggered rapid GTP hydrolysis (Fig. 2A, left). Similarly, the addition of the PH domain alone did not induce GTP hydrolysis, but subsequent addition of the isolated ZA domain initiated rapid GTP hydrolysis (Fig. 2A, middle). A plausible hypothesis is that the PH domain is the Arf binding site, possibly contributing to catalysis by affecting Arf allosterically. We corroborated the results using LUVs for membrane surface and an assay measuring GTP hydrolysis directly. Including the PH domain in the reaction increased activity over that observed with ZA alone, with ZA in the presence of the PH domain having a half-maximal effect at approximately 4.10^-6^ M and inducing complete or near complete GTP hydrolysis at saturating concentrations (Fig. 2B). The ^ΔN14^PH domain, which binds PI(4,5)P_2_ as efficiently as the *wt* PH domain, had 1/100^th^ the effect of the *wt* PH domain, consistent with the N-terminal extension of the PH domain contributing to the allosteric mechanism or alternatively, increasing affinity of the PH domain for Arf.

**Fig. 2.**
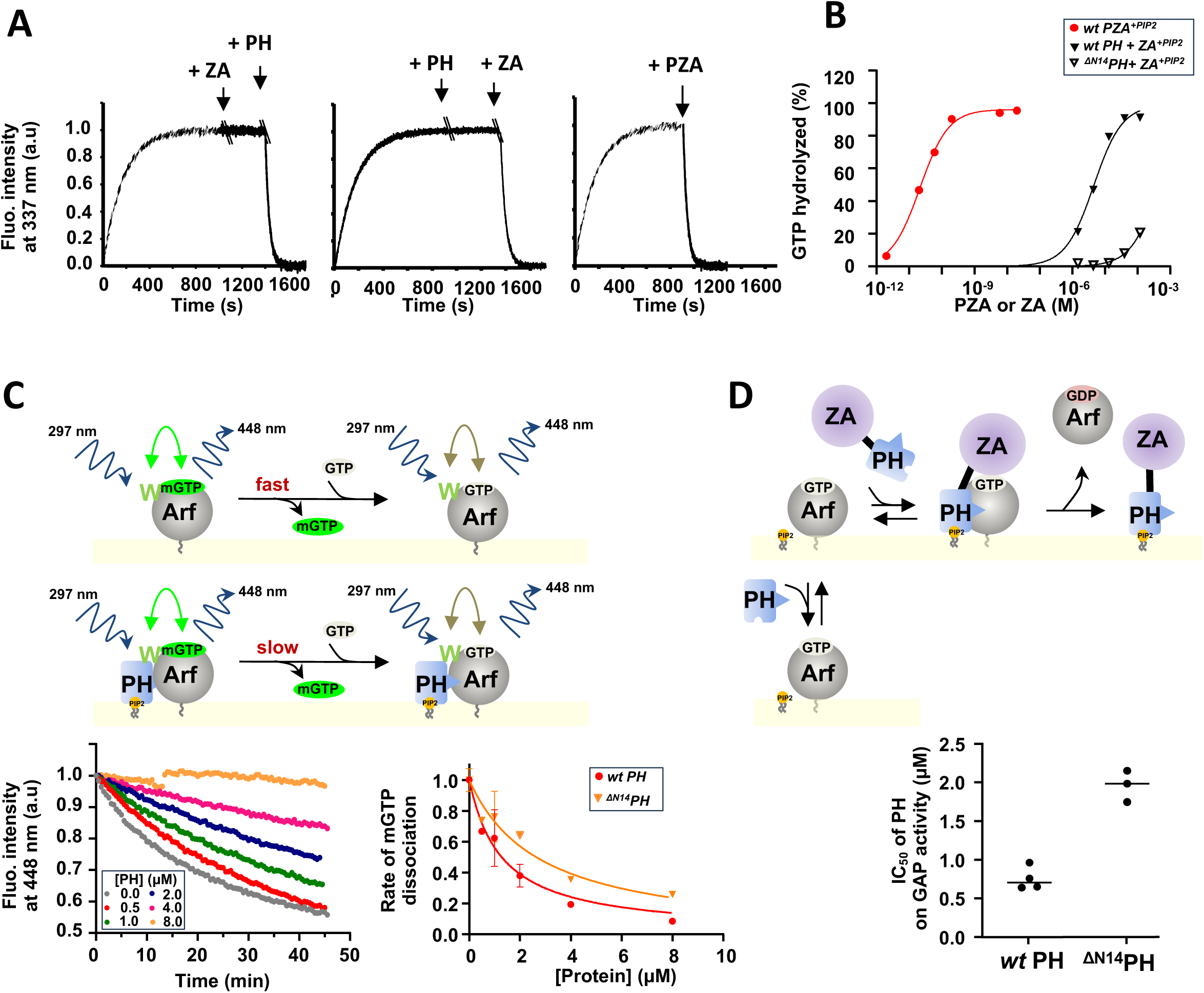
Binding of ASAP1 PH allosterically prompts myrArf1 for GTP hydrolysis. **A)** Representative tryptophan fluorescence kinetics trace of GTP hydrolysis of myr-Arf1 after GDP/GTP exchange triggered by the addition of PH and ArfGAP-Ankyrin repeat domains either isolated or in tandem. **Left:** Isolated PH domain (5μM) was added to myrArf1·GTP (5μM) bound to ND for ∼ 500 s before addition of ZA domain (25 μM). **Middle:** Isolated ZA domain (25 μM) was added to myrArf1·GTP (5μM) for ∼ 600 s before addition of PH domain (5μM). **Right:** *wt* PZA domain (10 nM) was added to myrArf1·GTP (5μM) when GDP/GTP exchange was complete (800 s). Nucleotide exchange Arf·GDP (5 μM) was triggered by the addition of 2 mM EDTA in the presence of 20 μM GTP, at 22 °C in the presence of ND containing 0.5 mM of accessible lipids. **B)** The percentage of GTP bound to myr-Arf1 hydrolyzed in 3 min is plotted against the concentration of ArfGAP-Ankyrin repeat domains after addition of ASAP1 PH-ArfGAP-Ankyrin repeat domains in tandem (●), of isolated wt-ASAP1 PH domain (▪) or of isolated ^ΔN14^ASAP1 PH (□). **E) Top:** Principle of FRET based assay used to measure K_d_ of PH for Arf1. MyrArf1 was loaded with the nucleotide analog mant-GTP, and the concentration of mant-GTP -bound Arf was followed by Fluorescence Resonance Energy Transfer (FRET) from myrArf1 tryptophan to mant-GTP. Then, *wt* PH or ^ΔN14^PH was titrated into the reaction containing an excess of GTP. Here, a decrease in FRET indicates mant-GTP dissociation. If the dissociation rate is slower for myrArf1·GTP in complex with the PH domain than for uncomplexed myrArf1·GTP, the concentration of PH domain reducing the rate to ½ the maximum reduction in rate is then equal to the dissociation constant (Kd). **Left:** Example of kinetic of FRET intensity measured at increasing concentration of PH domain. **Right:** Dissociation rates as a function of PH domain concentration. The dissociation rates are normalized to the rate measured in the absence of PH domain (0.05 min^-1^) **F) Top:** Principle of GAP activity assay used to measure K_d_ of PH for Arf1. **Bottom:** Concentration of PH domain necessary to reduce by half GAP activity (IC50) in reactions containing myrArf1·GTP at concentrations less than the K_m_ for *wt* PZA (1.10^-6^ M) and 1.10^-9^ M *wt* PZA. Under these conditions, half-maximal inhibition occurs at approximately the K_d_. IC50 values (the concentration of PH domain required to reduce GTP hydrolysis by ½ in 3 min) from each independent experiment are shown. Error bars represent standard deviation. ****, p < 0.0001 via one-way ANOVA with repeated measures (and mixed effects) and Dunnett’s multiple comparisons test against WT.

To discriminate between the two, we monitored the rate of GTP dissociation and GAP activity of *wt* PZA in the presence of *wt* PH or ^ΔN14^PH domain. These assays served as proxies for their respective affinity for myrArf1 (Fig. 2C, D). We found that the ^ΔN14^PH domain had ∼ half the affinity of the *wt* PH domain for Arf·GTP. Given that ^ΔN14^PZA had less than 1/100^th^ of the GAP activity of *wt* PZA and the ^ΔN14^PH domain less than 1/100^th^ of the “*in trans”* effect of *wt* PH, the data support that the N-terminal extension of the PH domain binds to and allosterically modifies Arf to facilitate GTP hydrolysis. Additionally, the K_d_ measured by both assays was similar to the K_m_ for *wt* PZA (1-2 μM)^3,22^, consistent with the hypothesis that the PH domain is the substrate binding site in ASAP1.

### Binding of ASAP1 PH to myrArf1 by NMR

To gain insight into how ASAP1 PH facilitates GTP hydrolysis, we observed NMR spectra of myrArf1 in complex with ASAP1 PH at the surface of negatively charged NDs containing the phosphoinositide PI(4,5)P_2_ using Transverse Optimized Relaxation Spectroscopy (TROSY) techniques^23^ (Fig. 3A, B).

**Fig. 3.**
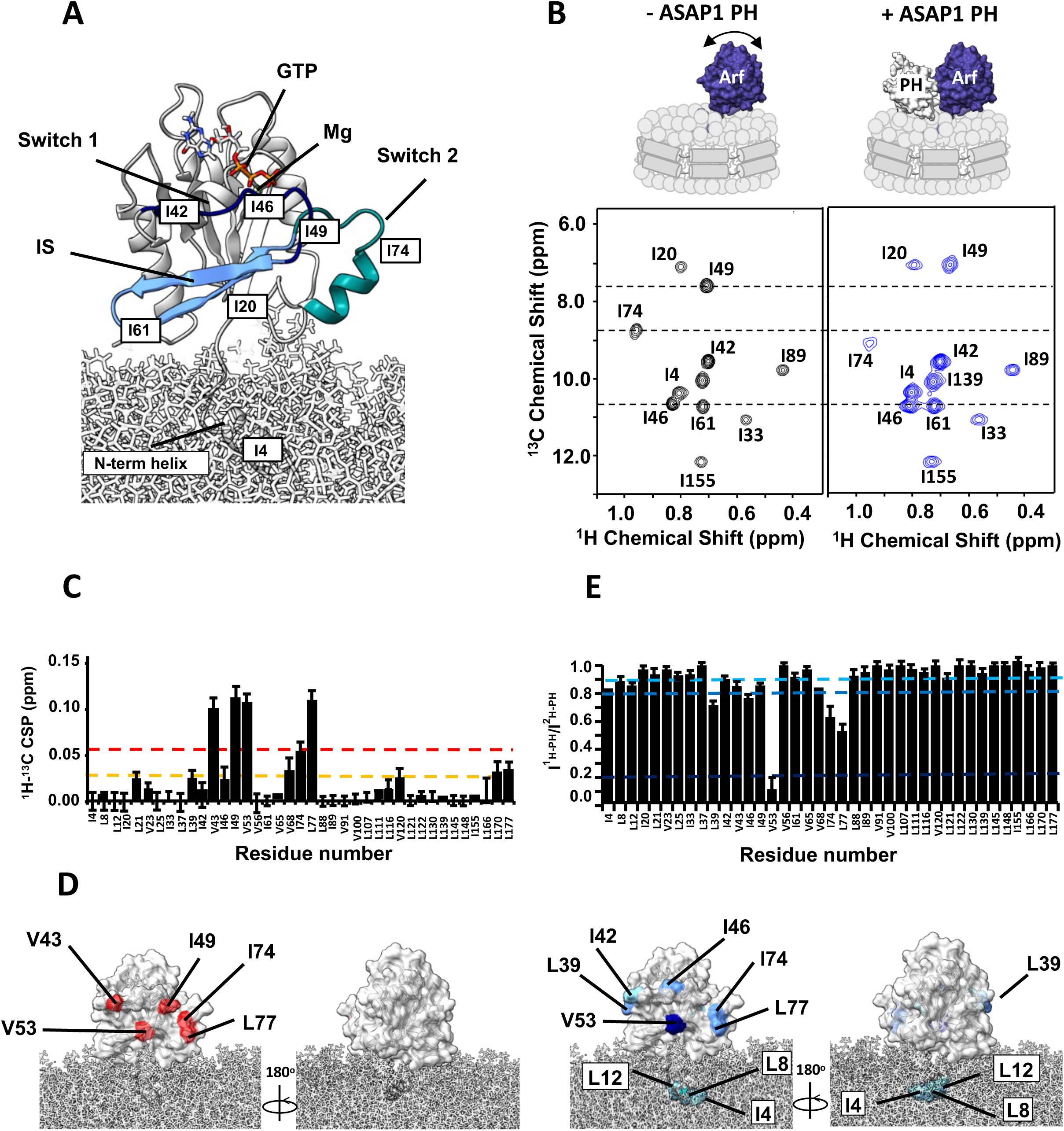
Binding of ASAP1 PH to myrArf1 at the membrane. **A)** Homology model of human myr-Arf1^31^ is shown in grey ribbon format at the membrane surface. Switch 1 (residues 40-49, dark blue), switch 2 (residues 68-78, cyan) and the interswitch (residues 50-67, light blue) are highlighted. Residues 2–13 form the N-terminal helix (embedded in the membrane), and residues 17–181 constitute the G-domain (solvent-exposed). Images created using Chimera^58^. **B)** ^1^H-^13^C HMQC spectrum centered on the Ile region of 50 µM U-^2^H,^15^N, δ1-^13^C^1^H-labeled Ile, δ1/δ2 - ^13^C^1^H-labeled Leu and γ1/γ2-^13^C^1^H-labeled Val myr-Arf1·GTP on a nanodisc (PC:PI(4,5)P_2_ = 95:5) in the absence (left, black) or in the presence of ^2^H ASAP1 PH (right, blue) at a Arf:PH ratio of 1:1.2) **C)** ^1^H-^13^C CSPs observed between myrArf1 free or bound to ASAP1 PH domain (Arf:PH 1:1.2) plotted against residue number. **D)** ^1^H-^13^C CSP values are mapped on the Arf1 surface: CSP> 2σ (red), CSP >1σ (orange). **E)** Ratio of intensities of Arf1 ^1^H-^13^C methyl cross peaks measured in the presence of ^1^H ASAP1 and ^2^H ASAP1 PH plotted against residue number. **F)** Ratios are mapped on the Arf1 surface with I^1H-PH^/I^2H-PH^ < 0.8 (dark blue) and 0.9 < I^1H-PH^/I^2H-PH^ < 0.8 (light blue). In addition to strong effects observed for V53 and residues of switch 1 and 2, the small amount of broadening observed for methyl groups on the N-terminal of myrArf1 may indicate proximity of the membrane bound part of myrArf1 with the ASAP1 PH domain as proposed in Roy et al^4^.

We first analyzed the spectral perturbations induced on a ^1^H-^13^C methyl Heteronuclear Multiple Quantum Coherence (HMQC) spectrum of δ1-^13^C^1^H-labeled isoleucine (Ile), δ1/δ2 - ^13^C^1^H-labeled Leucine (Leu) and γ1/γ2-^13^C^1^H-labeled Valine (Val) and otherwise perdeuterated myrArf1·GTPγS in the presence of an equimolar ratio of uniformly ^2^H (U-^2^H) labeled ASAP1 PH at the surface of ND (Fig. SI1). In addition to the uniform broadening of myrArf1 resonances due to complex formation, specific chemical shift perturbations (CSPs) and selective resonance attenuation were seen on switch 1 (Val43, Ile49), switch 2 (Ile 74, Leu77) and the interswitch region (Val53) (Fig. 3C and Fig. SI2). Those perturbations form a well-defined patch on one side of Arf1, suggesting that those residues are part of the interface with ASAP1 PH (Fig. 3D). Some smaller but significant changes were also seen on Leu 170 in C’ helix, which might result from indirect coupling effects between the two lobes of the G domain, and Leu 177 in the C’ helix and Val120 at the membrane facing tip of the β_6_ strand, which might result from new interactions of those residues with lipids.

Chemical shift mapping data were complemented by comparing intensities of myrArf1 methyls in the presence of protonated (I^1H^) or perdeuterated (I^2H^) ASAP1 PH as in^24^ (Fig. 3E). A low I^1H^/I^2H^ indicates proximity between a methyl group and ASAP1 PH. The methyl resonance of interswitch residue Val53 broadened beyond detection when bound to protonated ASAP1 PH but was well resolved when bound to deuterated ASAP1 PH, indicating that Val53 is buried at the interface, in agreement with chemical shift mapping data. Smaller effects were observed for switch 1 (Ile42, Val43, Ile46 and Ile 49) and switch 2 residues (Ile74, Leu77), suggesting that those methyls are either only transiently part of the interface or not as deeply buried as Val53 (Fig. 3F).

### Binding of myrArf1 to ASAP1 PH by NMR

To determine the surface of the PH domain that might be in the Arf1:PH interface, we monitored spectral perturbations on isotopically labeled *wt* PH when bound to an equimolar ratio of ND-associated myrArf1 in a ^1^H-^13^C methyl HMQC spectrum. The *wt* PH domain was expressed as U-^2^H,^15^N and δ1-^13^C^1^H-labeled Ile, δ1 -^13^C^1^H-labeled Leu and γ1-^13^C^1^H-labeled Val, β -^13^C^1^H-labeled Alanine (Ala) and γ2-^13^C^1^H-labeled Threonine (Thr). The largest CSPs were observed on strands β_5_ (Thr387, Cys388, Gln389, Val390), β_6_ (Leu402) and β_7_ (Thr408) and the C-terminal alpha helix capping the β sandwich (Ile423, Leu434, Thr435 and Ala437) (Fig. 4A). Those perturbations form well-defined patches on one side of ASAP1 PH suggesting that strands β_5-7_ and the capping alpha helix are part of the interface with myrArf1 (Fig. 4B). Smaller CSPs on the other side of the PH domain i.e., at the tips of strands β_2_ and β_3_ (Val364, Ile368), suggests that a small fraction of *wt* PH might have a different orientation when interacting with myrArf1 (Fig. 4B).

**Fig. 4.**
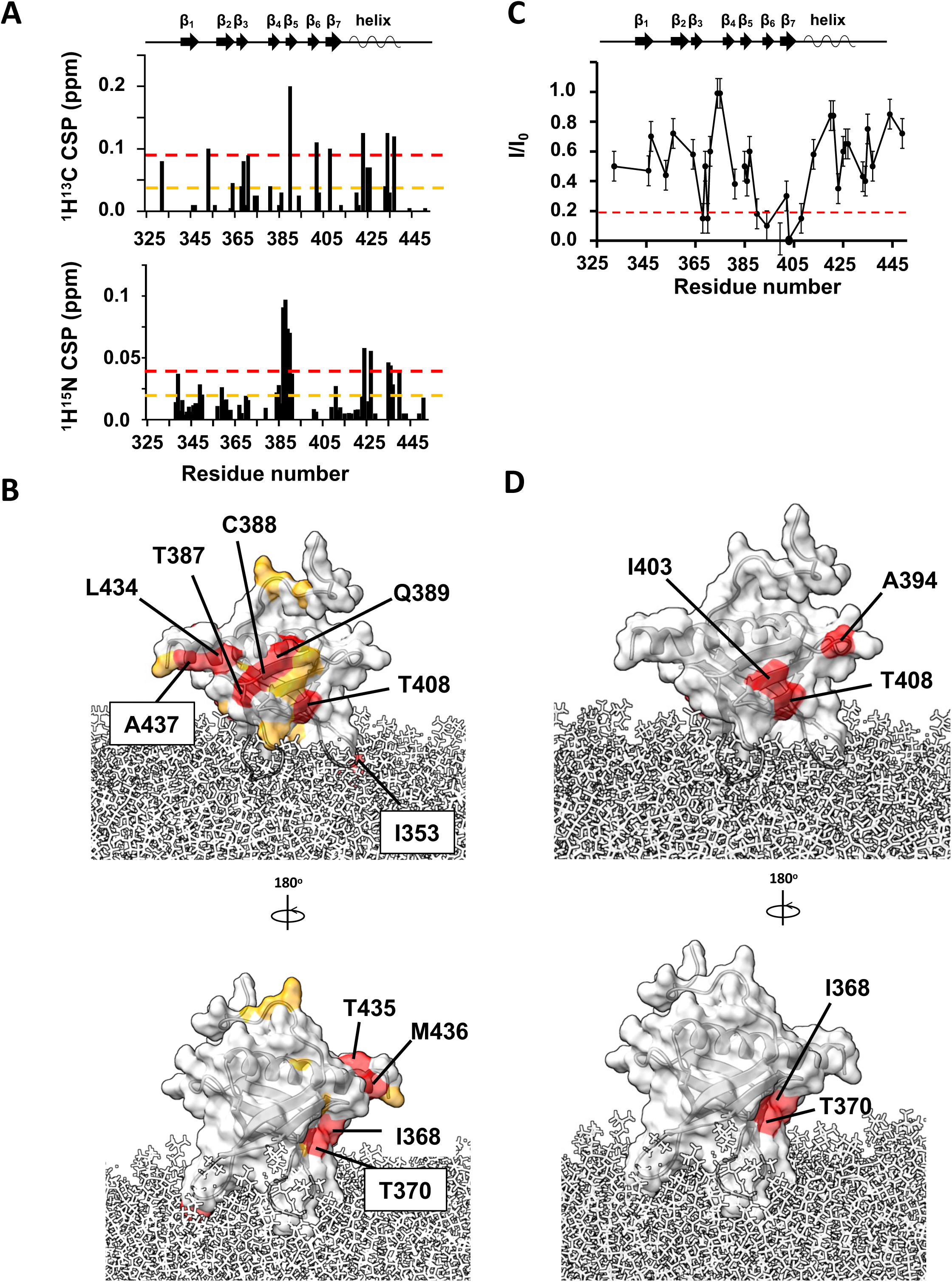
Binding of myrArf1 to ASAP1 PH at the membrane. **A)** ^1^H-^13^C (top) and ^1^H-^15^N CSPs (bottom) observed between ASAP1 PH bound to ND only or ND in the presence of myr-Arf1 (PH:Arf1 1:1.2) plotted against residue number. **B)** CSP values are mapped on the Arf1 surface: CSP > 2σ (red), CSP >1σ (orange). In addition to the large CSPs observed on the β sandwich, the large CSP observed for Ile353 (β_1_/β_2_ loop) could result from a reorientation of *wt* PH at the membrane in the presence of myrArf1 or from a direct interaction with myrArf1. Two independent experiments were performed. Data are presented as mean values. Error bars were calculated based on the signal-to-noise (S/N) ratio of the spectra as described in Methods. **D)** PRE <0.2 in the presence of MTSL-tagged myrArf1K38C mapped on the ASAP1 structure. Two patches were observed on opposite side of the β sandwich: The first patch includes residues Val390 (β_5_ strand), Ala394 (β_5_/β_6_ loop), Leu402, Ile403 ((β_6_ strand) and Ala413 on β_7_ strand (called thereafter β_5_/β_7_ patch). The second patch includes residues Ile368 and Thr370 on the β_3_ strand (called thereafter β_2_/β_3_ patch).

Intermolecular paramagnetic relaxation enhancement^25,26^ (PRE) effects induced on ASAP1 PH by spin-labeled myrArf1 variants were used to determine distance restraints between the interacting partners. To that end, two myrArf1 variants were obtained, in which Arf1 native cysteine (Cys159) was mutated to Ala and Lys38 (near switch 1), or Lys73 (near switch 2) were replaced with Cys to allow for spin-labeling. Their functional interaction with ASAP1 was largely unchanged compared to myrArf1 (Fig. SI3). A nitroxide spin label (MTSL) was covalently linked to each of the two myrArf1 variants via a disulfide bond and used to measure PREs to methyl groups of ASAP1 PH. Ratios of intensities of ASAP1 PH resonances in the presence of each of the two spin-labeled variants in their paramagnetic (I, intensity with spin label) and diamagnetic (I_0_, intensity with reduced spin label) states are shown in Fig. 4C and Fig. SI4. Mapping the residues with I/I_0_ < 0.2 (corresponding to distance < 16 Å from the paramagnetic site location) onto the surface of ASAP1 PH (Fig. 4D) shows they localized within two patches (called thereafter β_5_/β_7_ patch and β_2_/β_3_ patch), on opposite sides of ASAP1 PH and overlapping with CSPs.

### Model of myrArf1:ASAP1 PH complex

A model of the myrArf1:ASAP1 PH complex was built based on NMR experimental restraints by docking using the HADDOCK suite^27,28^. Because the two patches identified by PREs are separated by more than 20Å, the observed PRE patterns cannot result from a single ensemble of similar orientations and be consistent with CSPs data, suggesting two states in dynamic equilibrium. Therefore, we chose to build 2 different sets of unambiguous distance restraints based on their location. Set^β_5_/β_7_^ and Set^β_2_/β_3_^ were built with PRE ratio < 0.2 observed on the β_5_/β_7_ patch and β_2_/β_3_ patch, respectively. Both sets were complemented by ambiguous interaction restraints based on CSPs (Table SI2). We utilized those restraints in docking calculations to obtain models of the myrArf1:ASAP1 PH complex (Clusters^β_5_;/β_7_^ and Clusters^β_2_;/β_3_^). We then used representative members of each cluster as starting conformations for multi-replicas, multi-μs long all-atoms MD simulations in the presence of PC: PI(4,5)P_2_ membranes (SIM^β_5_/β_7_^ SIM^β_2_/β_3_^). ^1^H-transverse relaxation for the *wt* PH domain methyls were calculated every ns and averaged over of the length of SIM^β_5_/β_7_^ and SIM^β_2_/β_3_^ . We then compared the population weighted back-calculated PRE to the experimental ones (see material and methods and SI). We found that back-calculated PREs correlated best with experimental PREs with a ratio β_5_/β_7_ : β_2_/β_3_ of ∼ 9:1 (slope of 1.03 and Pearson’s coefficient of 0.95), indicating that the interface predominantly involves the state wherein the β_5_/β_7_ side of ASAP1 PH interacts with myrArf1 (Fig. 5A, B) with a buried surface area of ∼ 1100 Å^2^ (Fig. 5C).

**Fig. 5.**
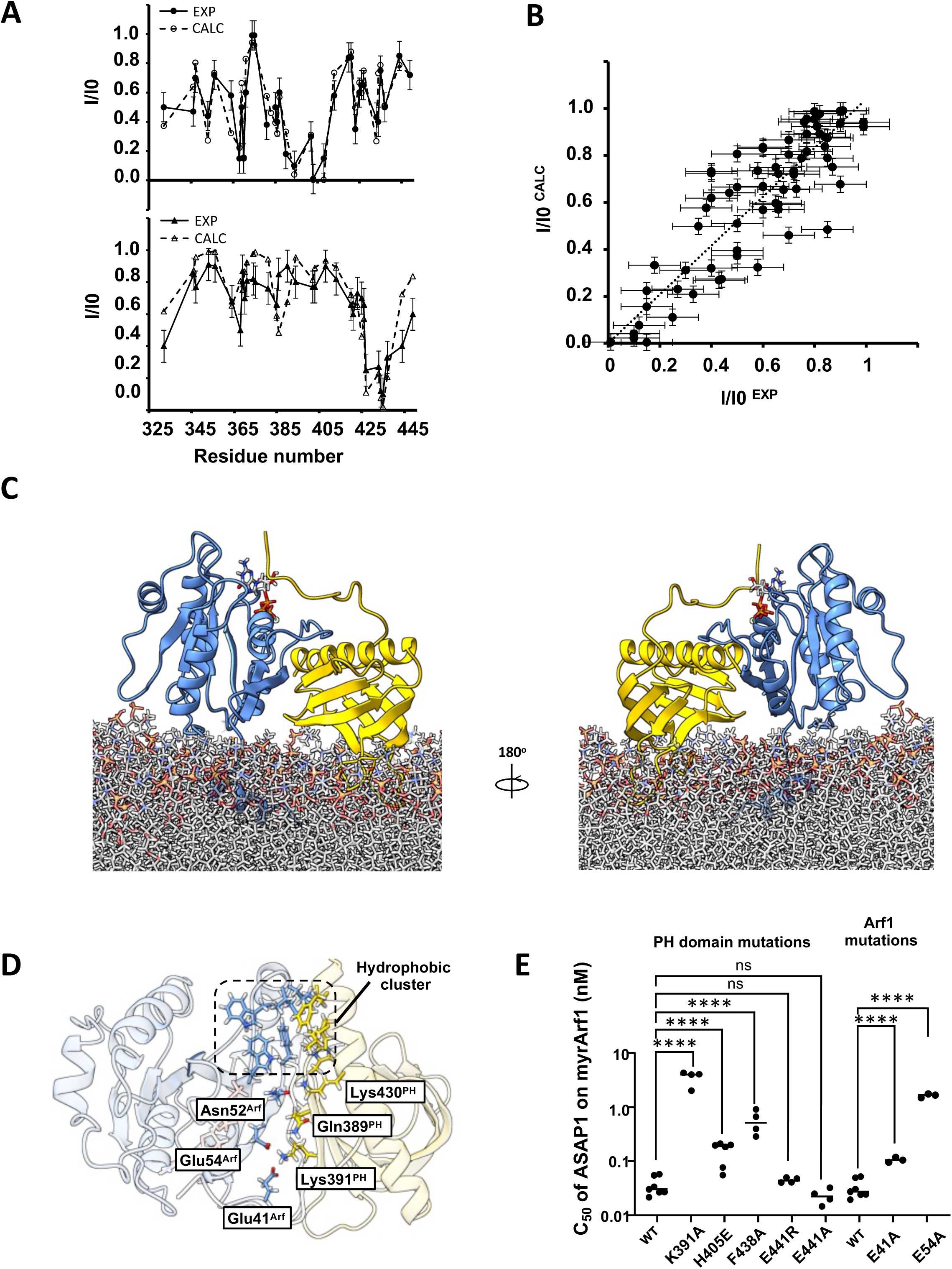
Model of ASAP1 PH:myrArf1 complex. **A)** Comparison of experimental (●) and calculated PRE (○) profile for ASAP1 PH in the presence of MTSL-tagged myrArf1K38C (top) and MTSL-tagged myrArf1K73C (bottom) **B)** Correlation of calculated PRE (y-axis) versus measured PRE (x-axis) corresponding to **A)**. For the measured PRE, error bars were calculated based on the spectral signal-to-noise ratio as described in Methods. For the calculated PRE (y-axis), data are presented as mean values over all MD simulations replicas +/−SD. **C)** Snapshot representative of the model of the complex between myrArf1 (blue ribbon) with the GTP nucleotide represented as ball and stick and ASAP1 PH (gold ribbon) at the membrane surface. **D)** Key interactions at the interface. Stable intermolecular salt bridges and residues forming the hydrophobic cluster comprising Phe51, Trp66, Ile74, Leu77 and Trp78 on Arf1 and Leu 386 (β_4_/β_5_ loop), His405 (β_6_/β_7_ loop) and Leu 434, Ala 437 and Phe438 (end of the C-terminal helix) on ASAP1 PH are shown. **E)** Effect on GAP activity assays of selected mutations on Arf1 (left) and ASAP1 PH (right). C50 values (the concentration of PZA required to achieve 50% of GTP hydrolysis in 3 min) from each independent experiment are shown. Error bars represent standard deviation. ns, not significant; ****, p < 0.0001 via one-way ANOVA with repeated measures (and mixed effects) and Dunnett’s multiple comparisons test against WT.

Seven intermolecular hydrogen bonds/salt bridges are consistently detected in the MD trajectories. Three involve Arf1 side chains, Glu41 (Switch 1) and Glu54 (interswitch (IS)), that form salt bridges with the sidechain of Lys391 (strand β_5_) on ASAP1 PH and Asn52 (IS) that forms transient salt bridges with either Lys430 (C-terminal helix) or Gln389 (strand β_5_) of the PH domain. Hydrogen bonds of the Phe51 and Val53 main chains (myrArf1) with Lys430 or Glu431 and Gln389 side chains of ASAP1 PH, respectively, provide additional stability. These interactions are supplemented by hydrophobic clusters between the IS and switch 2 regions of Arf1 and the β_4_/β_5_, β_6_/β_7_ loops and the end of the C-terminal helix of ASAP1 PH. Interestingly, transient hydrogens bonds are also detected between residues of the N-terminal extension of ASAP1 PH (Gly325, Thr327, Met329, His330, Gln331) and switch 1 of myrArf1 (Thr44, Thr45 and Ile46). Key interfacial interactions are illustrated in Fig.5D. To validate the structural model of myrArf1: ASAP1 PH complex, we examined the effect of mutations on binding and GAP activity (Fig. 5E and Table SI3-5). For example, mutating Lys391 on ASAP1 PH or Glu54 on myrArf1 to Ala reduced ArfGAP activity by 100- and 70-fold while mutating His405 or Phe438 on ASAP1 PH to Glu or Ala lowered it by 5- and 15-fold (Fig. 5E). These results are consistent with the prediction that these residues play a role at the interface. Similarly, the decrease in GAP activity for the Phe438 mutation is expected from a reduction in sidechain hydrophobicity in the hydrophobic cluster. Mutating Glu441 to Ala/Arg had no effect on ArfGAP activity, which is consistent with the model predicting that this residue is not part of the interface. Interestingly, mutating Lys391 to Ala only reduced binding by 2-fold, as expected from the loss of free energy upon removing a Lys-Glu salt bridge^29^ (Fig. SI5). Given that this mutation had a disproportionate effect on catalysis, similar to that observed upon removing residues 325-338 of the *wt* PH domain, it suggests that those residues are critical to the allosteric mechanism.

### The ZA domain has a minimal contribution to wt PZA binding to myrArf1·GTP

We also tested the hypothesis that binding of the PH domain to Arf could affect the energetics of ZA binding. A Cys159-MTSL tagged myrArf1 was used to measure intermolecular PREs between myrArf1 (bound to a non-hydrolysable GTP analog), either alone or in complex with *wt* PH, and the isolated, isotopically labeled ZA domain, at the membrane surface of NDs. The absence of intermolecular PREs (100μM of Arf or Arf: PH and 100 μM of ZA domain) indicated that less than 5% of the ZA domain was bound to Arf under those conditions, consistent with a K_d_ for ZA·Arf in the 0.1-1.0 millimolar range and little or no effect of *wt* PH on the K_d_ (Fig. SI6). This was complemented by monitoring ^1^H-^15^N NMR signals in TROSY-HSQC^23^ of domain segmentally labeled PZA^30^ (only the ZA domain was isotopically labeled) bound to membrane-bound myrArf1 and conducted at 25°C. Contrary to Arf for which the orientational space is reduced when bound to the PH domain^31^, the conformational space explored by the ZA domain is not reduced in the presence of the Arf: PH complex, as would have been expected from a high affinity complex between ZA and Arf1 (Fig. 6A, B). Taken together, this shows that binding of the PH domain to Arf minimally changes, if any, the weak affinity of the ZA domain for Arf.

**Fig. 6.**
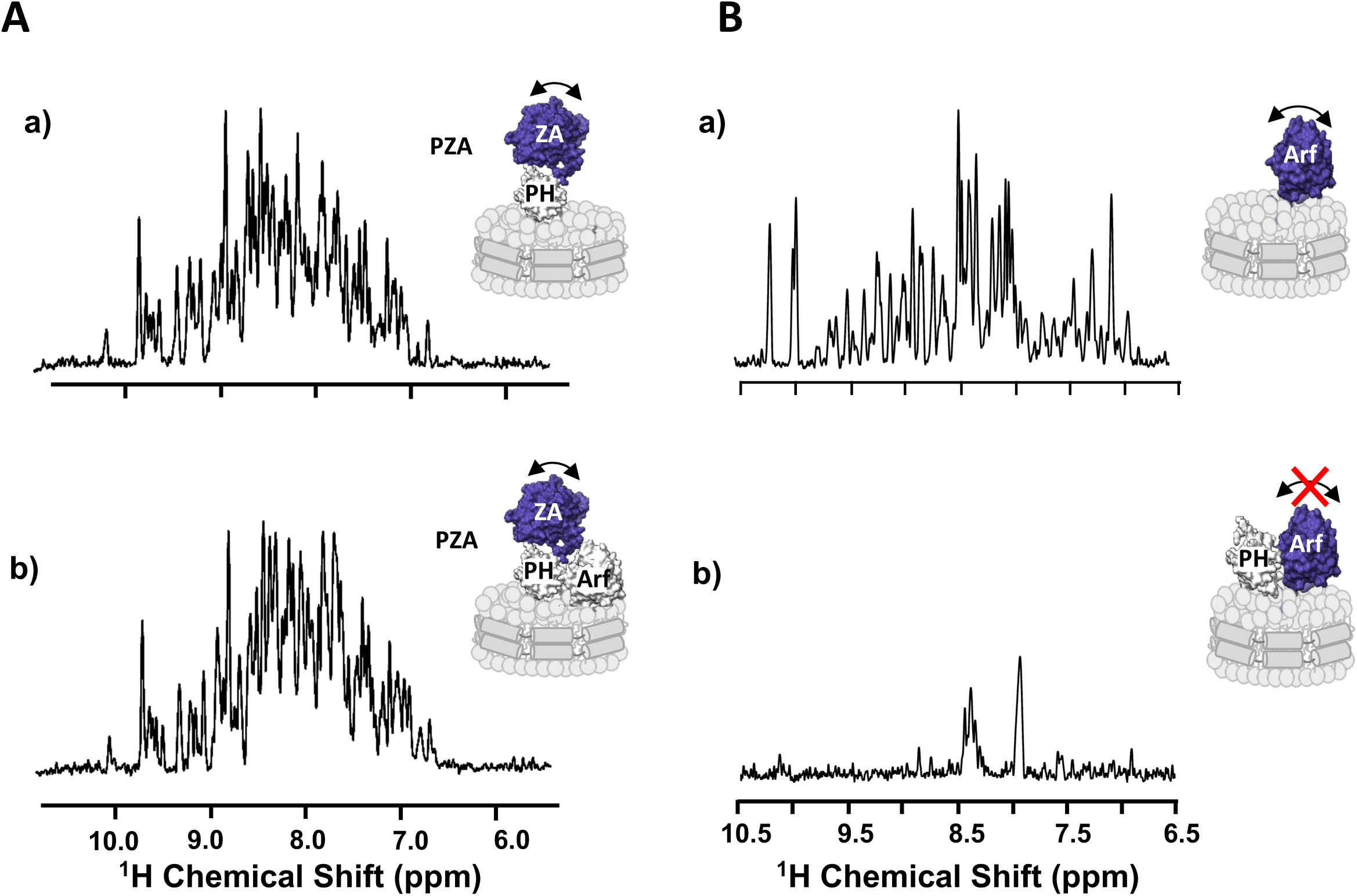
The ZA domain binds weakly to the Arf:PH complex. **A)** One-dimensional ^1^H NMR spectrum corresponding to the projection of an ^1^H-^15^N TROSY-HSQC along the F1 dimension obtained for ZA domain labeled PZA bound to ND alone (a) or bound to Arf (b), measured at the membrane surface of NDs containing PI(4,5)P_2_. The absence of visible changes indicates that the ZA domain reorients independently from the complex between Arf, PH and the ND, as expected from weak affinity between ZA and the ARF:PH complex. Motional reorientation is represented by a double-sided arrow. Isotopically labeled, NMR visible domain used in each experiment is colored in blue. **B)** One-dimensional ^1^H NMR spectrum corresponding to the projection of an ^1^H-^15^N TROSY-HSQC along the F1 dimension obtained for myrArf1 bound to ND alone (a), in the presence of *wt* PH (b).The addition of a stoichiometric ratio of ASAP1-PH resulted in the loss of nearly all amide backbone resonances in the ^1^H-^15^N TROSY-HSQC spectrum, in stark contrast with the spectrum in the absence of the PH domain indicating that the G domain becomes locked, reorienting with the same correlation time as the nanodisc. Motional reorientation is represented by a double-sided arrow. Isotopically labeled, NMR visible domain used in each experiment is colored in blue.

### Binding of the PH domain to Arf remodels the GTP binding pocket toward a transition state intermediate

MD simulations were used to examine the hypothesis that the PH domain might stabilize a transition state intermediate towards catalysis. Complex formation destabilized the internal hydrogen bond (H-bond) network of myrArf1 leading to (i) an outward motion of residues 41-45 of switch 1 and (ii) a subtle rearrangement of polar residues in the active site. In the simulations, Glu54^Arf1^ was a pivotal residue connecting key functional segments of the small GTPase. Without the PH domain, Glu54^Arf1^ forms a salt bridge with Lys38^Arf1^ and interacts with Thr44^Arf1^. When Glu54^Arf1^ binds to PH, it forms a stable salt bridge with Lys391^PH^. Consequently, Glu54^Arf1^ participates less in intramolecular hydrogen bonding causing an outward motion of residues 41-45 (Fig. 7A, B). Simulations suggest that the N-terminal extension of the PH domain stabilizes this conformation. Inferences from the MD simulations were tested using NMR. Altering the length of the N-terminal extension of ASAP1 PH specifically affects Val43 and Ile46 of switch 1, indicating proximity between residues 325-334 of *wt* PH and switch 1 of Arf (Fig. 7C) in agreement with the simulations. The spectral perturbation for Val43 was abolished by either randomizing the N-terminal extension amino acid sequence or mutating Tyr327 to Ala but was only reduced by mutating His330 and/or Gln331 to Ala (Fig. SI7). This indicated that (i) the tyrosine ring caused a ring current effect that altered the chemical shifts of Val43, demonstrating proximity between the two, and (ii) the interaction with switch 1 is sequence dependent. We also found that switch 1 residues were less affected by protonation/deuteration when bound to ^ΔN14^PH than to *wt* PH (Fig. 7D), consistent with the loss of transient contacts between the N-terminus and switch 1 when myrArf1 is complexed to a shortened version of the PH domain.

**Fig. 7.**
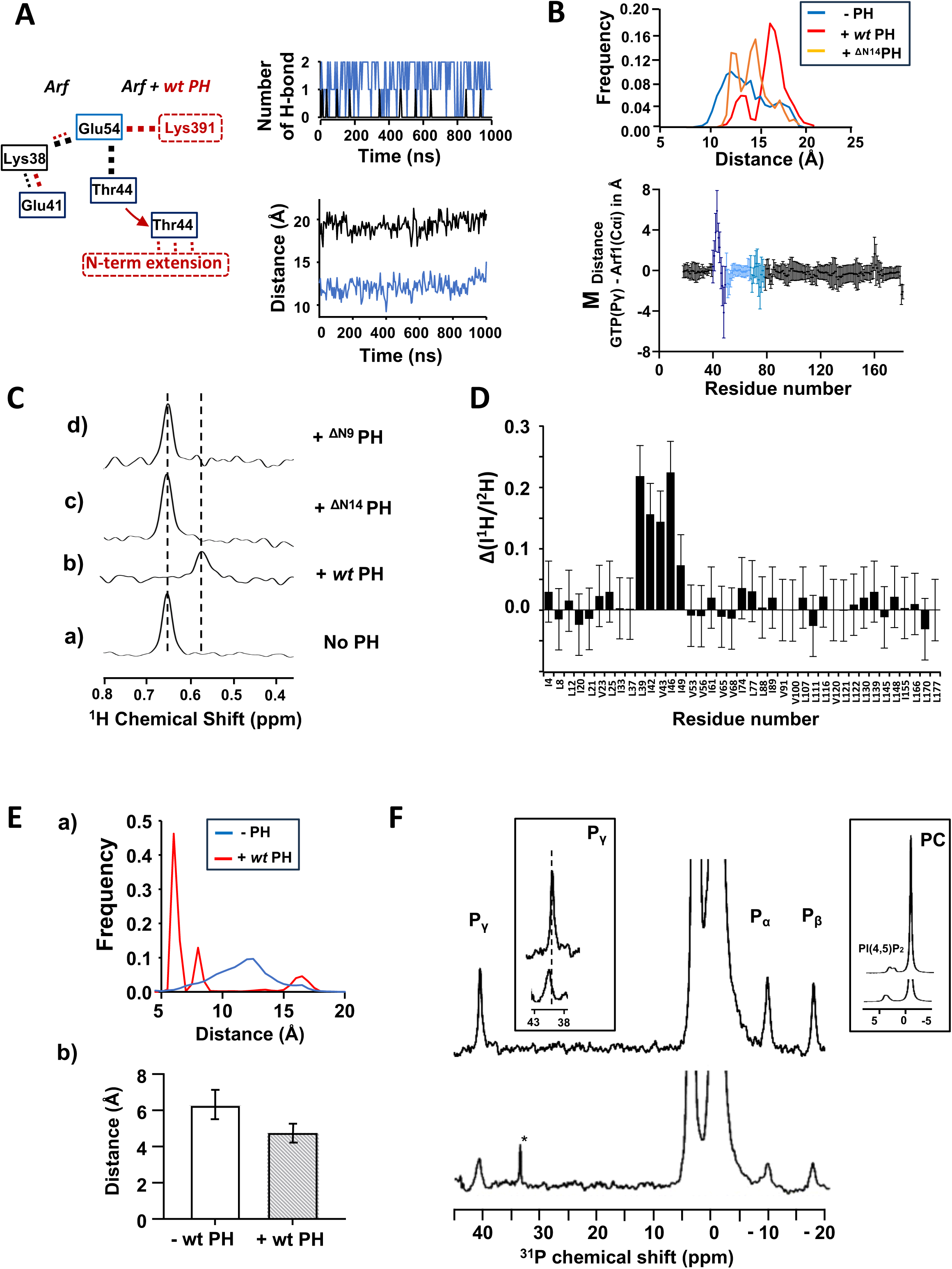
The PH domain destabilizes the internal Hydrogen bond network of Arf1. **A)** (a) Changes in the hydrogen bond network linking Glu54, Thr44 and Lys38 in Arf1 upon interaction with *wt* PH (b) Time dependence of the number of H-bonds formed between Glu54 and Thr44 for Arf alone (blue) or for Arf in complex with *wt* PH (black). Only one of the replicas is shown as an example. (c) Time dependence of the distance between the center of mass (COM) of residue 39-45 of switch 1 and GTP-Pγ corresponding to the replicas shown in (b) **B) (top)** Histogram of the distance distribution between COM of residue 39-45 of switch 1 and GTP-Pγ for Arf, Arf + *wt* PH and Arf + ^ΔN14^PH calculated over the entire length of the simulations **(bottom)** Plot of the average distance difference calculated between GTP-Pγ atom of GTP and carbon alpha (CA) of each Arf1 residue calculated as (d^GTP-Pγ→CAi^(Arf:*wt* PH) - d^GTP-Pγ→CAi^(Arf)). Error bars are calculated as the sum of the SD. (**C)** Stack of rows extracted from a ^1^H-^13^C HMQC experiment along the proton dimension of Val43 (myrArf1) in the absence (a) or in the presence of *wt*-ASAP1 PH (b), ^ΔN14^ASAP1 PH (c) or ^ΔN9^ASAP1 PH (d). **D)** Difference ΔI between the ratio of intensities of Arf1 ^1^H-^13^C methyl cross peaks measured in the presence of ^1^H ASAP1 PH and ^2^H ASAP1 PH using wt-ASAP1 PH or ^ΔN14^ASAP1 PH plotted against residue number. ΔI is calculated as (I^1H-wt-PH^/I^2H-wt-PH^)/ (I^1H-ΔN14-PH^/I^2H-ΔN14-PH^). **(E) (top)** Histogram of the distance distribution between CA of residue K30 and GTP-Pγ for Arf (blue) and Arf + *wt* PH (red) calculated over the entire length of the simulations **(bottom)** Plot of the average distance calculated between Mg^2+^ and atom OD1 of Asp 67 for Arf (left) and Arf + *wt* PH (right). **(F)** ^31^P NMR spectra of Arf bound to GTPγS in the presence (bottom) and absence of ASAP1 PH (top). Left inset shows deshielding of GTP-Pγ chemical shift in the presence of ASAP1 PH. Right inset shows deshielding of PI(4,5)P_2_ phosphate groups due to PH binding. *, trace γ-P_i_.

There were also subtle movements of polar residues in the active site of Arf. Residues 28-30 of the P loop are found on average, closer to the γ-phosphate of GTP. This is seen in 4 out of 5 replicas (Fig. 7E) and leads to an increase in the likelihood of the sidechain NH_3_-group of Lys30 to establish an H-bond with the γ-phosphate oxygens (Fig. 7E). This might participate in the stabilization of the transition state during GTP hydrolysis^32–34^. Because several changes involve residues near the nucleotide-binding site, we examined the effects on the bound nucleotide that accompanied complex formation using naturally occurring ^31^P resonances of GTPγS bound to myrArf1 alone or in complex with ASAP1 PH. As observed previously^35^, only one set of resonance lines is observable at 25°C indicating that myrArf1·GTPγS exists predominantly in a single (NMR-distinguishable) conformational state at the membrane (Fig. 7F, top). The addition of the *wt* PH domain triggers (i) a shift of the γ-phosphate resonance line indicating a change in its electronic environment upon PH complexation (Fig. 7F, bottom, left inset) (ii) an increase in line width at half height of all phosphate lines as expected from complexation (Fig. 7F, bottom) and (iii) a shift of the PI(4,5)P_2_ phosphate lines attributed to the binding of the PH domain to PI(4,5)P_2_ headgroups (Fig. 7F, bottom, right inset). These data suggest that contacts between ASAP1 PH and myrArf1 alter the conformation, dynamics, and solvation of functionally relevant regions around the nucleotide binding pocket compatible with the Arf:PH complex at the membrane surface being a pre-transition state intermediate.

### The PH domain increases the catalytic rate of the GTPase reaction

Based on the results presented, we concluded that the ASAP1 PH domain affects the rate of GTP hydrolysis by three mechanisms: (i) membrane recruitment (ii) Arf1 binding; and (iii) allosteric activation, whereby the PH domain induces a conformational change in Arf. To quantify the relative contribution of each mechanism, we modeled the reactions of PH, ZA and Arf1 for the *“in trans”* with a system of 11 nonlinear ordinary differential equations (ODEs) and the *“in tandem”* with a system of 11 ODEs (see SI for details)^36–38^. A simultaneous fit to the data (Fig. 8A and Fig. 8B-E) indicated that: (i) while dimensional reduction and substrate binding can each increase GAP activity by roughly three orders of magnitude compared to ZA alone, the combination of these two effects enhances GAP activity by only four to five orders of magnitude as catalysis becomes the rate-limiting step and (ii) the remaining three of the roughly eight orders of magnitude increase in GAP activity between ZA alone and *wt* PZA in bilayers containing PI(4,5)P_2_, can be attributed to a PH-triggered allosteric change in Arf1 causing a 1000-fold increase in the catalytic rate k_cat_ of the reaction (Table SI6). This agrees with conformational changes located around the nucleotide binding site of myrArf1 observed in simulations and by NMR. The change in k_cat_ only arises through the simultaneous association of the PH domain with PI(4,5)P_2_ and Arf1 at the membrane. One model prediction represents a hypothetical scenario in which PI(4,5)P_2_ function is limited to membrane localization (Fig. 8A); that is, PI(4,5)P_2_ binding to the PH domain does not affect the k_cat_ or the rate constant for association of the ZA domain to Arf. This result combines the enhancement effects of membrane association and of tethering ZA to PH. As can be seen, the combined effects do not fully account for the observed GAP activity of *wt* PZA with 5 mol% PI(4,5)P_2_, excluding this simplified model. The remaining three orders of magnitude increase in GAP activity arise through a 1000-fold increase in k_cat_ upon PH_m_ (PH bound to PI(4,5)P_2_) binding to Arf. In addition, halving the affinity of the PH domain for Arf (simulating the difference in affinity between ^ΔN14^PZA and *wt* PZA) in a model with recruitment and binding, but without conformational change in Arf resulted in a near negligible impact on overall GAP activity by PZA. These results are consistent with the critical role of the intrinsically disordered region of the PH domain for GAP activity.

**Fig. 8.**
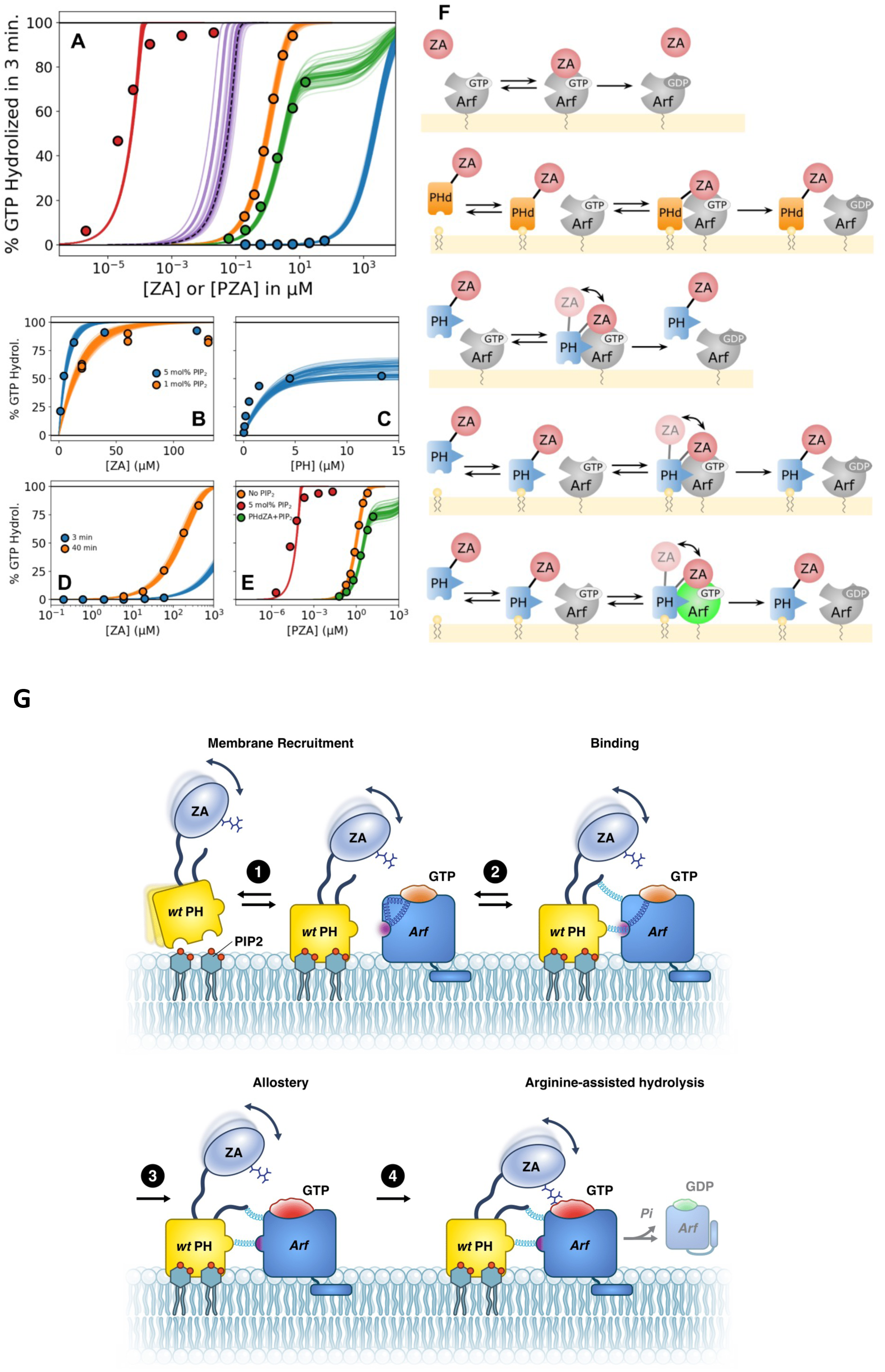
Kinetic modeling of GAP activity. (A) Fraction of GTP hydrolyzed in 3 minutes by either ZA or PZA. From right to left, curves are for ZA (blue), PHdZA with PI(4,5)P_2_ (green), PZA (orange), hypothetical PZA with PIP_2_ which only provides localization (dashed purple), and PZA with PI(4,5)P_2_ (red). Points represent experimental data and lines are predictions from optimized kinetic ODE model. In all cases where PI(4,5)P_2_ is present its concentration is 5 mol%. (B–E) Plots of experimental data (circles, triangles) used to optimize the kinetic model, along with model predictions (solid curves) for the final optimal parameter set. Initial Arf·GTP concentration is 1μM in all cases. Unless indicated otherwise, PI(4,5)P_2_ concentration is 5 mol%. Reaction time is 3 minutes except where otherwise indicated. Data and model curves in D (3 min) and E are the same as those shown in A. (F) Schematic diagrams of the dominant pathways for each scenario plotted in A. Top to bottom correspond to curves from right to left in A. Enhancement of Arf GAP activity through direct interaction with PH is indicated by purple color change of Arf, direct interaction with PH_m_ indicated in vibrant green. The final step of each line is catalyzed GTP hydrolysis. (G) PH domain as an integral element of a catalytic interface in the Arf GAP ASAP1.Step 1: The PH domain serves as a phosphoinositide binding module to translocate ASAP1 PZA to the membrane. Step 2: The PH domain acts as the primary binding site for its substrate Arf·GTP, the physical link to the Arf GAP domain providing proximity to the catalytic arginine in the Arf GAP domain. Step 3: The formation of the complex alters Arf intramolecular H-bond network and triggers the remodeling of the nucleotide binding site in Arf·GTP to achieve a transition state intermediate. Step 4: a catalytic arginine in the Arf GAP domain of ASAP1 facilitate the nucleophilic attack of water on GTP.

## Discussion

We defined the interface between myrArf1 and the ASAP1 PH domain on the membrane, allowing us to determine the structural basis and mechanistic underpinnings for enhancement of catalytic activity. The results indicate that the PH domain accelerates hydrolysis of GTP bound to Arf at the membrane surface via three distinct mechanisms (Fig. 8G). First, under the *in vitro* assay conditions, the PH domain serves as a phosphoinositide binding module to translocate ASAP1 PZA to the membrane where it searches a restricted volume for its substrate, increasing collision frequency and thereby decreasing time to collision. Second, the PH domain acts as the primary binding site for its substrate Arf·GTP, the physical link to the Arf GAP domain providing proximity to the catalytic arginine in the Arf GAP domain and DxxGQ glutamine in the switch 2 region of Arf1. Third, the formation of the complex alters an Arf intramolecular H-bond network and triggers the remodeling of the nucleotide binding site in Arf·GTP to lower the free energy barrier for formation of the transition state intermediate. For ASAP1 PZA, the contribution of substrate binding and remodeling of the GTP binding pocket are of similar magnitude and together are greater than the contribution of membrane recruitment. In this model, the PH domain is an integral component of the catalytic interface. PH domains in other Arf GAPs as well as other regulators of the Ras superfamily might function similarly. The results are relevant to Arf signaling in that, at least in this case, the GAP domain is not a complete GAP with the substrate binding site residing in a separate domain. Thus, the substrate for the GAP domain is not Arf but Arf bound to another domain. The new conceptual model of PH domain function we propose is distinct from previous models in that the PH domain is an integral component of the catalytic interface, serving as both the primary substrate binding site and contributing directly to changes in the substrate necessary for efficient catalytic activity.

The interface between Arf and the PH domain is greater than 1000 Å^2^ and, when bound to Arf1, the ASAP1 PH domain disrupts the intramolecular H-bond network that occurs in the small GTPase. Experiments and computations suggest essential roles of internal H-bond networks in shaping protein conformational dynamics^39,40^. For instance, in G-Protein Coupled Receptors, an extensive hydrogen bond network that spans all functional motifs of the protein in the active state has been identified, suggesting it might contribute to their activation mechanism. Here, the PH domain establishes competing salt bridges with Arf residues that are critical to the organization of the active site. The result is an intermediate with some, but not all characteristics of the transition intermediates observed in crystal structures of members of the Ras superfamily in complex with their GAP proteins^41^. In particular, we did not observe major changes in the conformational space explored by Arf1 Gln71 (Gln61 in RAS), a key catalytic residue in the RAS superfamily^42,43^. It is likely that the GAP domain, which is absent here, would affect Gln71 conformation as shown for ASAP3^44^.

The interface between ASAP1 PH and Arf1·GTP differs from the three other Arf-PH domain complexes reported in the literature. These structures include crystal structures of a complex between N-terminally truncated Arf1 or Arf6 and the Arf binding domain of ARHGAP21^45^ or [247-399]Grp1^46^, and an NMR derived model of the interaction between yeast Arf1 and the PH domain of four-phosphate-adaptor protein 1 (FAPP1) at the membrane^47^. When overlaying Arf1 from our model to Arf1 in the FAPP1, ARHGAP21 or Grp1 complexes we observed that although all PH domains engage with a similar surface of Arf1, our model is the only one where strands β_5/7_ of PH interact with the interswitch of Arf1 at the membrane.

The paradigm shifting mechanism we propose provides a rationale for developing inhibitors for proteins with PH domains occurring in tandem with a catalytic domain. When the prevailing paradigm for PH domain function was membrane recruitment, efforts to develop inhibitors focused on blocking phosphoinositide binding to prevent recruitment, which has been described for Akt^48–52^. However, inhibitors targeting the PH domain of the Rho GEF pREX and Brag2 inhibit activity without blocking membrane association^53,54^. NAV-2729 binds to the PH domain of ASAP1, near one PI(4,5)P_2_ binding site^55^. Neither PI(4,5)P_2_ binding, nor membrane association are affected but GAP activity is inhibited. We speculate that the inhibitory small molecules affecting pREX and Arf exchange factors and GAPs might alter a conformation in the PH domain that is necessary for catalytic activity. Additional structural characterization and biochemical studies will be valuable for defining the inhibitory mechanisms.

The catalytic mechanism we defined here is important for understanding Arf biochemistry and biology. Our results indicate that the GAP domain of ASAP1 is not a complete GAP in that it lacks a substrate binding site. In ASAP1, the complete enzyme includes the PH and Arf GAP domains. Other Arf GAPs with PH domains N-terminal to the GAP domain may function similarly, but there are no obvious Arf binding sites in Arf GAPs without PH domains N-terminal to the Arf GAP domain. One plausible explanation is that the Arf binding site resides on another protein. This could explain why little or no in vitro activity has been detected for some of these Arf GAPs, e.g. Arf GAP2^56^, Arf GAP3, ADAP1 and AGFG1 (unpublished). Arf GAP1 activity is increased by including coatomer, a coat complex that binds to Arf1-GTP, in the reaction. Arf GAP2 has detectable activity in the presence of coatomer. These Arf GAPs might recognize Arf bound to effectors, and further, Arf GAPs might be considered subunits of effectors. In this way, Arf GAPs might be integral to effector function, consistent with the observation that in budding yeast, Arf GAPs can rescue Arf insufficiency.

In summary, the PH domain of ASAP1 is an integral element of the catalytic GAP site, which is a newly identified function for PH domains and essential for understanding the regulation and function of Arfs.

## Materials and methods

### Protein expression and purification

#### Expression and purification of myrArf1 and mutants

The human Arf1 (Uniprot P84077) gene was cloned into MCS1 of the pETDuet-1 vector (Novagen) between *Nco l* and *Sac I* restriction sites. A GSGSHHHHHH-tag was added at the C-terminus of human Arf1. The yeast NMT (Uniprot P14743) gene was subcloned into MCS2 of the pETDuet-1 vector between *Ndel I* and *Xho I* restriction sites.

For expression of the myr-Arf1 protein (∼23 kDa) for NMR studies, the pETDuet-1 plasmid was transformed into *E. coli* BL21 Star (DE3) cells (Invitrogen), plated on LB agar plate containing carbenicillin (100 mg/L) for overnight growth. For expression of non-isotopically labeled protein, the freshly transformed colonies were picked, resuspended into 50 mL of Luria-Bertani medium containing carbenicillin (Goldbio, C-103-50) and incubated in a shaking incubator at 37°C until an OD_600_ of about 0.6. The culture was then diluted one-to-one with 50 mL of fresh Luria-Bertani medium. After repeating the same dilution procedure twice, culture was then diluted one-to-four with 800 mL of fresh Luria-Bertani medium and incubated until an OD_600_ of about 1

before being diluted one to one and moved to a shaking incubator pre equilibrated at 22 °C (for a final culture volume of 2L). For the production of myr-Arf1, sodium myristate (Sigma-Aldrich, M8005) was added 10 minutes before induction to a final concentration of 100 μM and 50 mg/L coenzyme A sodium salt (Sigma-Aldrich, C3144) to promote efficient N-myristoylation. Protein expression was induced at an OD_600_ of 0.8 by adding isopropyl-β-D-thiogalactopyranoside (IPTG, Goldbio, I248C50) to a final concentration of 0.2 mM. For expression of U-[^2^H, ^15^N], ^13^CH_3_ methyl labeled protein, the freshly transformed colonies were picked and resuspended into 10 mL of M9/H_2_O media for overnight growth at 37°C in a shaking incubator. Then the overnight culture was poured into a fresh M9/H_2_O media with a total volume of 5 mL and OD_600_ of 0.2 and continued to grow until an OD_600_ of about 0.6. The culture was diluted one-to-one with M9/D_2_O medium (prepared with D-[^2^H;^12^C]-glucose, Cambridge Isotope Laboratories, Inc; DLM-2062-10 and ^15^N ammonium chloride, Sigma-Aldrich, 299251) and incubated until OD_600_ reached 0.6. After repeating the same dilution procedure twice (with the final culture volume of 40 mL), the cells were spun down and resuspended in 200 mL M9/D_2_O medium for overnight growth at 37°C^59^. The expression culture was made from the overnight culture by diluting to a volume of 2 liters with a starting OD_600_ of about 0.2. After the culture had reached an OD_600_ of 0.6, the temperature was reduced from 37°C to 22°C. For the production of myr-Arf1, sodium myristate (Sigma-Aldrich, M8005) was added 10 minutes before induction to a final concentration of 100 μM. At the same time, the media was supplemented with: 1) 50 mg/L 2-keto-3-[D_2_],4-[^13^C]-butyrate (Cambridge Isotope Laboratories, Inc. CDLM-7318) and 100 mg/L 2-keto-3-[D]-[^13^CH_3_,^12^CD_3_]- isovalerate (Cambridge Isotope Laboratories, Inc. CDLM-7317) to enable selective labeling of ILV methyl groups; 2) 50 mg/L coenzyme A sodium salt (Sigma-Aldrich, C3144) to promote efficient N-myristoylation. Protein expression was induced at an OD_600_ of 0.8 by adding IPTG to a final concentration of 0.2 mM. The culture was incubated for additional 16 hours at 22°C for protein expression.

Cells were harvested by centrifugation at 7,000 *g*, 4°C for 30 minutes. The cell pellets were resuspended in 25 mL lysis buffer (20 mM Tris-HCl, pH 8.0, 150 mM NaCl, 20 mM imidazole, 1 mM MgCl_2_, and 0.5 mM tris(2-carboxyethyl) phosphine (TCEP) with one tablet of EDTA-free protease inhibitor (Thermo Scientist, A32965). The cells were lysed with a model 110S microfluidizer (Microfluidics) and clarified by centrifugation at 48,000 *g* and 4°C for 45 minutes. The lysate was loaded onto two 5 mL HisTrap HP columns (GE Healthcare). After the columns were washed with six column volumes (CVs) of lysis buffer, Arf1 and myr-Arf1 were eluted with an identical buffer containing 300 mM imidazole in a linear gradient from 20 mM to 300 mM imidazole over 14 CVs. The purity of myr-Arf1 was examined by LC-MS. The fractions containing purified myr-Arf1 were pooled and kept at 4°C for further processing. The fractions containing both Arf1 and myr-Arf1 were combined and concentrated to a volume of one milliliter. Sodium chloride crystals were added to the sample to a final concentration of 3 M. After centrifugation at 21,000 × *g* and 4°C for 15 minutes, the supernatant was collected and applied to a 5 mL pre-equilibrated HiTrap Phenyl HP hydrophobic interaction column (GE Healthcare) using a running buffer with 20 mM Tris-HCl, pH 7.4, 3 M NaCl, 1 mM MgCl_2_, and 0.5 mM TCEP. After the column was washed with ten column volumes of running buffer, myr-Arf1 was eluted with 20 mM Tris-HCl (pH 7.4), 1 mM MgCl_2_, and 0.5 mM TCEP using a linear gradient. The purity of myr-Arf1 was confirmed by LC-MS. Combining with previously purified myr-Arf1, the myr-Arf1 was exchanged to a buffer condition of 20 mM Tris-HCl, pH 7.4, 150 mM NaCl, 1 mM MgCl_2_, and 0.5 mM TCEP, concentrated to about 100 μM, and stored at -80 °C for further usages. The protein concentration was calculated by measuring the absorbance at 280 nm using a molar extinction coefficient of 29,450 M^-1^ cm^-1^.

Mutants of myrArf1 were generated by site-directed mutagenesis of the template plasmid (human Arf1 and yeast NMT in pETDuet-1, described above) using custom DNA oligos (Integrated DNA Technologies). Briefly, for each individual reaction 25 ng of template DNA was used with 125 ng of each custom oligo to perform 18 cycles of mutagenesis using a high-fidelity DNA polymerase (PfuTurbo, Agilent). Afterward, the mutagenesis reactions were subjected to *DpnI* treatment (New England Biolabs) to remove non-mutated DNA. Each transformation reaction was then transformed in NEB® 5-alpha competent *E. coli* (New England Biolabs) according to manufacturer’s protocol and plated onto LB/Carbenicillin plates. Individual colonies were grown up in liquid media supplemented with Carbenicillin, after which the cell pellets were miniprepped (Qiagen) to isolate plasmid DNA. Clones were verified by Sanger sequencing with the pET Upstream primer (Novagen) at the NIH Center for Cancer Research (CCR) Genomics Core at the National Cancer Institute, Bethesda, MD. Colonies with desired Arf1 mutations (and no changes in yeast NMT) were then transformed into BL-21 Star (DE3) *E. coli* and used for expression and purification as listed above.

For expression of myArf1 and mutant proteins for binding and kinetic studies, the pETDuet-1 plasmid was transformed into *E. coli* BL21 (DE3) cells (Invitrogen), plated on LB agar plate containing ampicillin (100 mg/L) for overnight growth. Subsequent steps were as described above.

### Preparation of wt ASAP1 PH domain and mutants

The sequence for ASAP1 PH domains, [325-451]–ASAP1, [334-451] –ASAP1 and [339-451]–ASAP1 (∼14 kDa), was cloned between Nde I and Bam HI restriction sites of the pET3a vector, which was then transformed into *Escherichia coli* BL21 Star (DE3) cells (Invitrogen) for protein expression, plated on LB agar containing carbenicillin, and incubated overnight (o/n). After, the seed cultures were used to inoculate large-scale (1 L) cultures, and the cultures were grown until the OD600 reached ∼0.6 – 0.8 at which point they were induced with 1 mM IPTG for three hours at 37°C. Following induction, the cells were harvested by centrifugation, supernatant removed, and frozen at -80°C until purification.

PH domains were purified by resuspending in buffer A (50 mM Tris pH 7.4, 150 mM NaCl) by using 30 mL buffer for 1 liter’s worth of cell pellet. A single protease inhibitor tablet (cOmplete EDTA-free, Roche) was used for each cell pellet. The cells were lysed using a cell disruptor (Microfluidics) and then ultracentrifuged (6000*g* for 30 min) at 4°C. Afterward, the supernatant containing PH domain was removed, and all subsequent purification steps were conducted at room temperature as it was observed that the PH domain precipitates when chilled. The supernatant was applied to a 5 mL HiTrap SP HP column (Millipore Sigma) pre-equilibrated with buffer A, washed with 10 column volumes (CVs) of buffer A, then eluted with a 6 CV linear gradient from buffer A to buffer B (50 mM Tris pH 7.4, 1 M NaCl). Eluates containing PH domain were pooled and then injected onto a ∼120 mL HiPrep 16/60 Sephacryl S-100 HR SEC column (Millipore Sigma) pre-equilibrated with buffer A supplemented with 1 mM TCEP. SEC eluates containing PH were then pooled, concentrated using 3,000 MWCO spin-concentrators (Amicon), and snap-frozen using liquid nitrogen. Concentration of PH domain was determined by ultraviolet (UV) spectroscopy (ε^280^ = 16,960 M−1 cm−1).

For the production of [U-^2^H], [U-^15^N]-methyl specifically labeled protein, NH_4_Cl is substituted by ammonium chloride (^15^N ≥ 99%), D-glucose is replaced by d-(^2^H, ^12^C)-glucose (^2^H ≥ 98%), and ^13^CH_3_-methyl specifically labeled precursors are added as described below. For a typical cell culture of 500 ml, a few freshly transformed colonies of BL21 Star (DE3) cells were picked to inoculate 5 ml of M9/H_2_O minimal media for o/n growth at 37°C in a shaking incubator (250 rpm). One milliliter of the o/n culture [typical optical density at 600 nm (OD600) ∼ 1.2] was then used to inoculate 4 ml of fresh M9/H_2_O medium to achieve a starting OD600 of 0.25. At OD600 ∼ 0.5, 5 ml of M9/D_2_O minimal media was added and cell growth continued until an OD600 of ∼0.5 is reached. Cells were diluted again by a factor of 2 and growth followed to OD600 ∼ 0.5. This cycle was repeated until a D_2_O/H_2_O ratio of 3:1 (20 ml total) is reached. Cells were then harvested by centrifugation (3000*g* for 30 min) and resuspended in 25 ml of M9/D_2_O, and growth was continued in a 100-ml baffled flask until an OD600 of 0.5 is reached, before an additional 25 ml of M9/D_2_O was added for o/n growth at 37°C. When the o/n OD600 was between 1.3 and 1.5, the o/n cell expression (50 ml) was added to 500 ml of M9/D_2_O and growth followed at 37°C, up to OD600 ∼ 0.6. For selective δ1-^13^C^1^H-labeled Ile, δ1 -^13^C^1^H-labeled Leu, γ1-^13^C^1^H-labeled Val, β -^13^C^1^H-labeled Ala and γ2-^13^C^1^H-labeled Thr labeling, the PLAM-AβIγ1LV_proS_Tγ kit was used (NMR-Bio). After the addition of the precursor according to the manufacturer’s protocol, cell growth continued until an OD600 of approximately 0.8 at 20°C is reached, at which time protein expression was induced with the addition of 1 mM IPTG. After induction, another 2 g/liter of D-(^2^H, ^12^C)-glucose was added, and the culture was grown o/n at 20°C. All subsequent steps were carried as described above.

For expression of ASAP1 PH and mutant proteins for binding and kinetic studies, the pET21 vector plasmid was transformed into *E. coli* BL21 (DE3) cells (Invitrogen), plated on LB agar plate containing ampicillin (100 mg/L) for overnight growth. Subsequent steps were as described above.

### Expression and purification of PZA and mutants

The PH, Arf GAP, and ankyrin repeats (PZA construct, residues 325 – 724) of ASAP1 were expressed and purified as previously described (PMID: 9819391). Briefly, PZA with a 10x N-terminal His tag was expressed by transforming into BL-21(DE3) competent cells, then picking individual colonies and growing using LB/ampicillin media until seed cultures were at an OD600 of ∼0.6. After, the seed cultures were used to inoculate large-scale (1 L) cultures, and the cultures were grown until the OD600 reached ∼0.6 – 0.8 at which point they were induced with 1 mM IPTG for three hours at 37°C. Following induction, the cells were harvested by centrifugation, supernatant removed, and frozen at -80°C until purification.

PZA was purified by resuspending in nickel buffer A (20 mM Tris pH 8, 500 mM NaCl, 20 mM imidazole) by using 30 mL buffer for 1 liter’s worth of cell pellet. A single protease inhibitor (cOmplete EDTA-free, Roche) was used for each cell pellet. The cells were lysed using a cell disruptor (Microfluidics) and then ultracentrifuged at >100,000 *g* for one hour at 4°C. Afterward, the supernatant containing PZA was removed. The supernatant was applied to a 1 mL HisTrap HP column (Millipore Sigma) pre-equilibrated with nickel buffer A, washed with 10 column volumes (CVs) of nickel buffer A, then eluted with a 10 CV linear gradient from nickel buffer A to nickel buffer B (20 mM Tris pH 8, 500 mM NaCl, 500 mM imidazole). Eluates containing PZA were pooled and then injected onto a ∼120 mL HiPrep 16/60 Sephacryl S-200 HR SEC column (Millipore Sigma) pre-equilibrated with storage buffer (20 mM Tris pH 8, 150 mM NaCl). SEC eluates containing PZA were then pooled, concentrated using 20,000 MWCO spin-concentrators (Amicon), and snap-frozen using liquid nitrogen.

Mutants of PZA were generated by site-directed mutagenesis of the template plasmid (PZA in pET-19b) using custom DNA oligos (Integrated DNA Technologies). Briefly, for each individual reaction 25 ng of template DNA was used with 125 ng of each custom oligo to perform 18 cycles of mutagenesis using a high-fidelity DNA polymerase (PfuTurbo, Agilent). Afterward, the mutagenesis reactions were subjected to DpnI treatment (New England Biolabs) to remove non-mutated DNA. Each transformation reaction was then transformed in NEB® 5-alpha competent E. coli (New England Biolabs) according to manufacturer’s protocol, and plated onto LB/ampicillin plates. Individual colonies were grown up in liquid media supplemented with ampicillin, after which the cell pellets were miniprepped (Qiagen) to isolate plasmid DNA. Clones were verified by Sanger sequencing with standard T7 forward and terminal primers at the NIH Center for Cancer Research (CCR) Genomics Core at the National Cancer Institute, Bethesda, MD. Colonies with desired PZA mutations were then transformed into BL-21(DE3) E. coli and used for expression and purification as listed above.

### Preparation of isotopically domain labeled PZA

Domain labeled PZA was obtained using Sortase mediated ligation (SML) and purified as previously described with some modifications^30^. Briefly, the reaction buffer (SML buffer) was 50 mM Tris, pH 7.8, 5 mM CaCl2, 100 mM NaCl, 10 mM L-Arginine, 0.2 mM TCEP. The stoichiometry for the reaction was ZA: PH: Srt 1:2:0.25. Typical reaction volume was 5 mL and the concentration of isotopically labeled ZA was set at 10 μM. The SML reaction was performed in a centrifugal concentrator [MWCO 3000 Da, Amicon] while spinning at 3000g. After a given time (10 min), the volume of the reaction was readjusted to 5 mL with SML buffer and 10μM of fresh PH domain was added. This was repeated 4 times for a total reaction of ∼ 45 min. Sortase ligated PZA was purified in a single step over a 5-ml SP anion exchange column (GE Healthcare) to minimize protein degradation. The column was equilibrated and washed with buffer A (20 mM Tris pH 7.4, 100 mM NaCl), and the protein was eluted using a 20-column volume gradient to 50% buffer B (buffer A and 1 M NaCl). The domain labeled PZA was usually eluted at 38 - 42 % buffer B. Buffer exchange into buffer A was done stepwise by successive centrifugation in a centrifugal concentrator [MWCO 3000 Da, Amicon] to minimize protein precipitation. The sample identity was confirmed by mass spectrometry, and purity was assessed by SDS– polyacrylamide gel electrophoresis. Concentration of domain labeled PZA domain was determined by ultraviolet (UV) spectroscopy (ε^280^ = 36900 M^−1^ cm^−1^).

### Expression and purification of MSPΔH5

The plasmid for MSPΔH5 was a gift from F. Hagn and G. Wagner (Harvard Medical School). The protein was expressed and purified as described previously^17^.

### Dissociation rate of mantGTP measured by Fluorescence spectroscopy

Binding and dissociation of mantGTP was monitored by following the FRET signal resulting from resonance energy transfer from Arf tryptophan to the methylanthronoyl (mant) group on GTP (excitation 290 nm; emission 448 nm) using a FluorMax3 spectrophotometer (Jobin Yvon Horiba, Edison, NJ). To load myrArf1 with mant-GTP (ThermoFisher, M12415), 1 mM myrArf1 was incubated at 30°C for 40-60 min in 25 mM HEPES, pH 7.4, 100 mM NaCl, 1 mM dithiothreitol, 0.5 mM MgCl_2_, 1 mM EDTA, 5 mM mant-GTP, and 0.5 mM Large Unilamellar Vesicles (LUVs) with 5% mol PI(4,5)P_2_. After Mant-GTP loading reached plateau, different concentrations of ASAP1 PH were added to the reaction mixture and excess GTP (160 μM) was added to initiate the mant-GTP dissociation from Arf. Arf·mantGTP has a FRET signal, whereas Arf·GTP does not; therefore, the exchange results in a decrease in fluorescent signal.

### Preparation of myr-Arf1·GTPγS anchored on MSPΔH5 nanodiscs for NMR spectroscopy

#### Preparation of empty nanodiscs

All lipids were purchased from Avanti Polar Lipids, Inc. To prepare nanodiscs, acyl chain perdeuterated 1,2-dimyristoyl-sn-glycero-3-phosphadidylcholine (DMPCd54) in chloroform solution (Sigma-Aldrich 319988) and 1,2-dioleoyl-*sn*-glycero-3-phospho-(1’-myo-inositol-4’,5’-bisphosphate) (PI(4,5)P_2_) in chloroform: methanol (Millipore MX0488): water (20:9:1) solution were mixed and air-dried with nitrogen flow before solubilization with cholate in aqueous buffer [20 mM tris-HCl (pH 7.4), 150 mM NaCl, and 75 mM sodium cholate (Millipore SX0420)]. Nanodiscs were assembled by mixing MSPΔH5 with solubilized lipids at a ratio of 1:45 (final cholate concentration of 18 mM) followed by the removal of cholate from the mixture with Bio-Beads SM2 resin (Bio-Rad, 152-8920), under o/n rocking at 22°. Assembled NDs were then purified via a Superdex-200 size exclusion column (GE Healthcare) and concentrated on a centrifugal concentrator [10 kDa molecular weight cutoff (MWCO), ThermoFisher Scientific]. The concentration of NDs was determined by UV spectroscopy (ε^280^ = 18,450 M−1 cm−1).

#### Preparation of myr-Arf1·GTPγS anchored nanodiscs

MyrArf1·GDP was incubated with freshly prepared nanodiscs in 20 mM Tris-HCl, pH 7.4, 150 mM NaCl, 0.5 mM MgCl_2_, 0.5 mM TCEP, 1mM EDTA, and 2 mM GTPγS (Millipore Sigma G8634). After incubation at room temperature for 30 minutes, myrArf1·GTPγS anchored on MSPΔH5 NDs was purified by a Superdex 200 Increase 10/300 GL column (GE Healthcare) into the final NMR buffer (20 mM Tris-HCl (pH 7.4), 150 mM NaCl, 1 mM MgCl_2_, and 0.5 mM TCEP). Typically, the fractions containing nanodisc-anchored myr-Arf1⦁GTPγS were pooled, concentrated and analyzed by SDS-PAGE. Known concentrations of Arf and MSP were run on the same SDS-PAGE gel and used to determine the Arf1: nanodiscs ratio. Concentration of NDs was then adjusted such that the ratio between Arf and ND was ∼ 1 by adding empty NDs.

#### Generation of spin-labeled variants of myrArf1

Spin-labeling conjugation reactions were performed on myrArf1·GTPγS anchored on nanodiscs purified in NMR buffer without reducing agent. The mono-cysteine variants of myrArf1 were treated with a 5-fold molar excess of S-(1-oxyl-2,2,5,5-tetramethyl-2,5-dihydro-1H-pyrrol-3-yl) methyl methanesulfonothioate (MTSL) (Toronto Research Chemicals Inc., 81213-52-7) and the reaction was allowed to proceed for 1 h at 25 °C in the dark. The unreacted spin-label was then removed by passing the protein samples through a Superdex 200 Increase 10/300 GL column pre-equilibrated in NMR buffer without reducing agent. The efficiency of the spin-labeling reactions was confirmed by LCMS mass spectrometry measurements. The reactions were found to proceed to completion in all cases.

### NMR spectroscopy

Experiments were performed using approximately 30-50 μM myrArf1(ASAP1 PH) anchored to 30-50 μM nanodiscs. Samples (approx. 250 μL) were contained in Shigemi microcells. Data were acquired at 25°C or 32.5 °C using Bruker AVIII-850, AVIII-800 and AVIII-700 spectrometers equipped with cryogenic TCI probes. All NMR data were processed and analyzed using Topspin and/or NMRPipe^60^.

^1^H,^13^C HMCQ spectra of ^13^CH3 methyl labeled proteins were acquired at 25°C using a SO-FAST HMQC pulse sequence as implemented in NMRlib package^61^. The spectral widths were set to 12.94 and 25 ppm in the ^1^H and ^13^C dimensions, respectively and inter-scan delays were set to 1 sec. In total, 1542 × 256 complex points were recorded, and between 16 and 64 scans/FID gave rise to an acquisition time between 1.5 and 5 hours. Prior to Fourier transformation, the data matrices were zero-filled to 4096 (^1^H) × 1024 (^13^C) complex points and multiplied by a cosine apodization function in both ^1^H and ^13^C dimensions.

^1^H,^15^N TROSY-HSQC spectra were acquired at 32.5°C using the BEST TROSY principle as implemented in NMRlib package^61^. The spectral widths were set to 12.06 and 36 ppm in the ^1^H and ^15^N dimensions, respectively with a recycle delay set to 1.5 sec. In total, 1368 × 192 complex points were recorded, and 96-128 scans/FID gave rise to an acquisition time between 5 and 7 hours. Prior to Fourier transformation, the data matrices were zero-filled to 4096 (^1^H) × 1024 (^15^N) complex points and multiplied by a cosine apodization function in both dimensions.

Chemical shift perturbations were calculated using Equation 1:

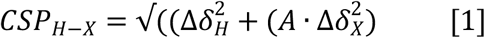

where Δδ_H_ is the change in amide proton value (in parts per million) and Δδ_X_ is the change in amide nitrogen or methyl carbon value (in parts per million). A is a scale factor equal to 0.17 (or 0.185) when X is ^15^N (or ^13^C).

1D-^31^P NMR spectra were acquired at 25 °C on a Bruker 700 MHz NMR spectrometer with a 5 mm Prodigy broadband cryogenic probe using 70° flip angle pulses, ∼ 15000 scans, an interscan delay of 7 s, an acquisition time of 84 ms, and a WALTZ-16 proton decoupling sequence.

#### PRE NMR measurements

Samples used for the paramagnetic relaxation enhancement (PRE) measurements contained 60 μM of protiated myrArf1 variants in their spin-labeled forms and 50 μM of U-^2^H,^15^N and δ1-^13^C^1^H-labeled Ile, δ1 -^13^C^1^H-labeled Leu and γ1-^13^C^1^H-labeled Val, β -^13^C^1^H-labeled Alanine (Ala) and γ2-^13^C^1^H-labeled Threonine (Thr) ASAP1 PH. For all spin-labeled variants, one sample each of the paramagnetic or the diamagnetic (in which the spin-label was reduced by incubation with 1 mM ascorbic acid for 2 h) species, were prepared. All measurements were performed at 25 °C. A recycle delay between scans of 4 s was used to insure adequate magnetization recovery for both the diamagnetic and paramagnetic states. The error values were calculated by the formula in equation (2):

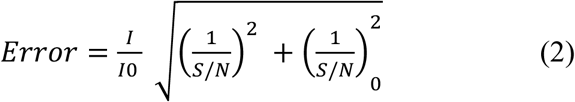

where *I*, (*S/N*) and I_0_, (*S/N*)_0_ are the intensity and signal-noise ratios of resonance measured in paramagnetic and diamagnetic samples, respectively.

### Docking

#### Generation of ambiguous interaction restraints

Ambiguous interaction restraints (AIRs) for use in the data-driven docking calculations (described below) were obtained for Arf1 and ASAP1 PH. Active residues were defined as those that were solvent exposed and displayed ^1^H-^13^C or ^1^H-^15^N CSPs > 0.1 ppm (i.e., > 2σ). Passive residues were defined as solvent exposed residues within 6.5 Å of the active set and/or with methyl (amide) CSPs between 0.047 (0.02) and 0.095 (0.04) ppm (i.e., 1σ < CSPs < 2σ) for *wt* PH and methyl (amide) CSPs between 0.03 (0.02) and 0.06 (0.04) ppm for Arf. Residues were defined as solvent exposed if they displayed at least 20% of relative solvent accessibility. The AIRs used for the docking calculations are listed in Table SI2.

#### Generation of unambiguous distance restraints from PREs

The intensity ratio of the HMQC spectra before (*I*) and after reduction with ascorbic acid (*I*_0_) were used to generate the unambiguous distance restraints. Protons with *I*/*I*_0_ < 0.2, including protons whose resonances were no longer detectable in the paramagnetic spectra, were assigned distance constraints ranging from 1.8 to 16 Å.

#### Data-driven docking

Structure calculations were performed using all ambiguous and unambiguous restraints using the HADDOCK suite^27,28^. The homology model of human myr-Arf1, generated based on yeast Arf1 as in, was used as the starting structure for Arf after truncating residues 1-17. The starting structure for PH was the AlphaFold model of the protein^62^. Docking simulations consisted of three consecutive stages. In the first, rigid body docking and energy minimization stage, a total of 10,000 structures of the complex were calculated allowing each of structures from the Arf1 and ASAP1 PH ensembles to explore a sufficiently broad landscape of initial orientations. At the end of this stage, 400 structures with the lowest energy scores were selected for the simulated annealing (second) and water refinement (third) stages. Residues 40 to 49 of Arf1 (switch 1) and 325-332 (N-terminal extension), 352-355 (β_1_/β_2_ loop), 374-378 (β_3_/β_4_ loop) and 445-451 (C_-_terminus) of ASAP1 PH were kept fully flexible during this stage of molecular docking. For Set^β5-β7^, 396 structures out of 400 were grouped into 1 major cluster. For Set^β2-β3^, 287 structures were grouped into 3 major clusters, each comprising of at least 20 individual structures, representing about 70% of the 400 water-refined models generated after the final refinement stage. The most relevant clusters for each set (Clusters^β_5_;/β_7_^ and Clusters^β_2_;/β_3_^), as defined by the most favorable HADDOCK score (HS, -128 ± 2.7 and -113.1 ± 0.8), contained ∼95 % (396 of 400) and ∼ 80% (213 out of 281) of the clustered structures, with an average RMSD value of 0.6 ± 0.6 Å and 2.6 ± 0.4 Å relative to the lowest energy structure.

### Molecular Dynamic Simulations

Because the initial docking stage was performed without membranes, we then used representative members of each cluster as starting conformations for all-atoms MD simulations.

Molecular systems for the starting point of simulations were built using CHARMM-GUI^63^. Relaxation using the standard steps provided by CHARMM-GUI was performed with NAMD. Production simulations employed the GPU optimized pmemd module of AMBER 18. The C36m forcefield was used for proteins^64^ and lipids^65^ with standard dynamics parameters (force-based switching between 10 and 12 Angstroms, SHAKE constraints on bonds to hydrogens, and the particle-mesh Ewald algorithm to handle long-ranged electrostatics with periodic boundary conditions). Zero tension at one atmosphere isotropic pressure was applied using a Monte Carlo barostat. Temperature was maintained at 310K with a Langevin friction coefficient of 1 ps^-1^. Five replicas of Arf, Arf:*wt* PH (Clusters^β_2_;/β_3_^) and Arf:^ΔN14^PH based on Arf:*wt* PH (Clusters^β_2_;/β_3_^) were run such that total simulation time for each system was at least two microseconds. Because we found that root mean square fluctuations (RMSF) between orientations over one replica was on the same order of magnitude as the RMSF between replicas for SIM ^β_5_/β_7_^, only one replica was run for Arf:*wt* PH Clusters^β_5_;/β_7_^ (SIM^β_2_/β_3_^).

The simulated lipid bilayer was composed of DMPC (ca. 295 lipids) and approximately 5% PIP2 (ca. 7 each of DOPI24 and DOPI25, the dioleoyl PIP2 in the C36 forcefield with varied protonation of the PIP2 phosphate). The total water layer was approximately 7.5 nm high (ca. 25000 TIP3P water molecules).

### Back calculation of PRE profiles from MD simulations

PREs were computed using the RotamerConvolveMD (version 1.3.2) package employing MDAnalysis (v1.0.0,^66^) as well as the rotamer library from^67^. The code was modified slightly to incorporate PREs using the target hydrogens, as opposed to the default (the modified code files are available at http://github.com/alexsodt/premod). Briefly, atomic coordinates of MTSL and of δ1-CH_3_-Ile, δ1/δ2-CH_3_-Leu and γ1/γ2-CH_3_-Val of Arf1 were extracted from MD simulations (1 frame every ns for each of the five multi μs long simulations for a total of ∼ 7500 frames) of Arf, Arf + wt PH and Arf + ^ΔN14^PH and used to back calculate PRE rates. For each of the ∼ 7500 frames, the effective PRE relaxation rate (*Γ*_2_^cal^) of a protein methyl proton was then calculated as the average of the PRE rate *Γ*_2,*i*_ computed for each of the *N* MTSL orientation the protein:

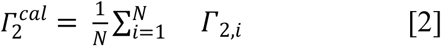

with *Γ*_2,*i*_ equal to:

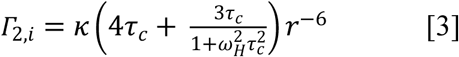

with *κ* equal to 1.23 × 10^-32^ cm^6^s^-2^ for the proton spin as reported previously, *r* the distance between the free electron and methyl group protons of Ile, Leu, Val, Ala or Thr residues in a single frame, *τ*_*c*_ the rotational correlation time of the electron-nuclear interaction, which was approximated using equation 4

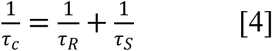

where *τ*_*R*_ is protein rotational correlation time and *τ*_*S*_ is the electronic longitudinal relaxation time. We used a *τ*_*R*_ of 70 ns for Arf in complex with PH and a value of 100 ns ^3^ for *τ*_*S*_. Calculation was then repeated for each frame and to back calculate average PRE rates for each methyl of SIM^β_5_/β_7_^ and SIM^β_2_/β_3_^

Fit to the data was then performed as a population weighted average of the PRE rate for each pose of the PH domain (Eq. 5).

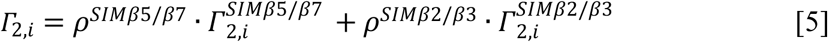

where *Γ*_2,*i*_ is, *ρ*^*SIMβ*_6_/*β*_7_^ and *ρ*^*SIMβ*_2_/*β*_3_^ are the population of SIM^β6/β7^ and SIM^β2/β3^, respectively. Then, the *Γ*_2_,*i* were converted to the intensity ratios of the paramagnetic (I) to diamagnetic (I_0_) peaks for representation 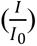 using equation 6:

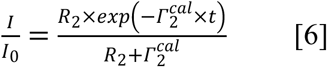

where *t* is the total evolution time of 6.89 ms in the HQMC pulse sequence, *R*_2_ is the intrinsic transverse relaxation rate, which was estimated from the half-height line width of peaks in the diamagnetic spectra.

### Functional assays

#### GDP-GTP exchange and GAP activity followed by Fluorescence spectroscopy on nanodiscs

All fluorescence experiments measurements were performed with a Horiba Fluoromax spectrofluorometer in a 120μL quartz cell. The sample (140μL) was thermostated at 22°C. The time constant of the fluorometer was set to 500 ms. The excitation wavelength (λ_exc_) and emission wavelength (λ_em_) were 297 and 337 nm, respectively. The excitation and emission bandwidth were set to 4 and 10 nm, respectively. Nucleotide exchange of purified myrArf1 (5μM) was assessed by monitoring the change in tryptophan fluorescence following addition of Ethylenediaminetetraacetic acid (EDTA) (2 mM) in the presence of 20 μM of GTP and 500 μM of exposed lipids in nanodiscs, which takes advantage of the large difference in fluorescence between the GDP- and GTP-bound forms of Arf proteins. The reaction was stopped by the addition of 1 mM MgCl_2_. Induction of hydrolysis of myrArf1·GTP to myrArf1·GDP was determined by following the change in tryptophan fluorescence, as previously described after addition of isolated ZA domain or *wt* PZA.

#### GAP activity followed by [α^32^P] radioactivity on LUV

LUVs were prepared by extrusion. Briefly, 1 µmol lipids (molar ratio, 40% PC, 25% PE, 15% PS, 10% cholesterol, and 10% total phosphoinositide) dissolved in chloroform, in a siliconized glass tube were dried under a nitrogen stream for 30 minutes to 1 hour, followed by lyophilization for at least one hour. The dried lipids were resuspended in 200 µL 1x PBS, for a final concentration of 5 mM. The solution was vortexed, subjected to five rounds of freeze/thaw, and extruded using a lipid extruder (Avanti Polar Lipids) through a Whatman Nucleopore Track-Etched membrane with 1 µM pores. The LUVs were stored at 4°C and were used within a week for activity assays.

GAP-induced conversion of myrArf1•GTP to myrArf1•GDP was determined as described previously. Reaction mixtures contained 25 mM HEPES, pH 7.4, 100 mM NaCl, 1 mM dithiothreitol, 2 mM MgCl_2_, 1 mM GTP, 0.5 mM LUVs, myrArf1 bound to [α^32^P]GTP, and variable concentrations of Arf GAP. The LUVs were included in the myrArf1 GTP loading reaction. The reactions were incubated at 30°C for 3 min. (unless otherwise specified), and quenched with 2 mL of ice-cold 20 mM Tris, pH 8.0, 100 mM NaCl, 10 mM MgCl2, and 1 mM dithiothreitol. Protein-bound nucleotide was trapped on nitrocellulose, and guanine nucleotide was released by addition of formic acid. [α^32^P]GDP and [α^32^P]GTP were then separated using thin-layer chromatography plates, and quantified.

### Mass Spectrometry Measurement

Mass spectra were obtained using Agilent Technologies 6100 Series Single Quadrupole LC/MS equipped with an electrospray source, operated in positive-ion mode. Separation was performed on a 300SB-C3 Poroshell column (2.1 mm × 75 mm; particle size 5 μm). The analytes were eluted at a flow rate of 1 ml/min with a 5 to 100% organic gradient over 5 min and holding the organic phase A for 1 min. Mobile phase A contained 5% acetic acid in water and mobile phase B was acetonitrile. Data acquisition, data analysis, and deconvolution of mass spectra were performed using Open Lab Chem Station Edition software (version C.01.05). Samples of purified proteins were typically 5 μl of a 5-10 μM solution.

### Statistical analysis

Data are expressed as means ±σ or ±SD when applicable. Representative data from two or more independent experiments were analyzed. P values were determined using the t-test.

## Data Availability

The data that support the findings of this study are available from the corresponding authors upon request.

## Acknowledgements

The research was supported by the Intramural Research Program of the National Cancer Institute (Projects ZIA BC 011419, ZIA BC 011131, and ZIA BC 011132 supported O.S., R.A.J., Y.Z., J.L., and R.A.B.; Project BC007365 supported X.J, E.M.R and P.A.R.) and by the Intramural Research Program of the Eunice Kennedy Shriver National Institute of Child Health and Human Development (Project 1ZIAHD008955 supported A.J.S). M.E.J. and S.J.F. gratefully acknowledge funding from an NIH MIRA Award R35GM133644 to M.E.J. F.H. was supported by the U.S. Department of Commerce, Award 70NANB17H299. The authors acknowledge the use of the Biophysics Resource, Center for Structural Biology, NCI, and the assistance of Dr. Sergey Tarasov and Ms. Marzena Dyba. The authors thank David Lambright, University of Massachusetts, for insightful discussions and review of the manuscript.

## Disclaimer

The content is solely the responsibility of the authors and does not represent the official views of the NIH. Certain commercial materials, equipment, and instruments are identified in this work to describe the experimental procedure as completely as possible. In no case does such identification imply a recommendation or endorsement by NIST, nor does it imply that the materials, equipment, or instrument identified are necessarily the best available for the purpose.

## Notes

### Competing Interest Statement

The authors have declared no competing interest.

## References

1. Kahn, R.A. et al. Consensus nomenclature for the human ArfGAP domain-containing proteins. Journal of Cell Biology 182, 1039–1044 (2008).

2. Sztul, E. et al. ARF GTPases and their GEFs and GAPs: concepts and challenges. Molecular Biology of the Cell 30, 1249–1271 (2019).

3. Jian, X.Y. et al. Autoinhibition of Arf GTPase-activating Protein Activity by the BAR Domain in ASAP1. Journal of Biological Chemistry 284, 1652–1663 (2009).

4. Luo, R. et al. Kinetic analysis of GTP hydrolysis catalysed by the Arf1-GTP-ASAP1 complex (vol 402, pg 439, 2007). Biochemical Journal 407, 471–471 (2007).

5. Lemmon, M.A. Membrane recognition by phospholipid-binding domains. Nat Rev Mol Cell Biol 9, 99–111 (2008).

6. Leonard, T.A., Loose, M. & Martens, S. The membrane surface as a platform that organizes cellular and biochemical processes. Dev Cell 58, 1315–1332 (2023).

7. Chan, T.O., Rittenhouse, S.E. & Tsichlis, P.N. AKT/PKB and other D3 phosphoinositide-regulated kinases: kinase activation by phosphoinositide-dependent phosphorylation. Annu Rev Biochem 68, 965–1014 (1999).

8. Das, S. et al. Structural Organization and Dynamics of Homodimeric Cytohesin Family Arf GTPase Exchange Factors in Solution and on Membranes. Structure 27, 1782–1797 e7 (2019).

9. DiNitto, J.P. et al. Structural basis and mechanism of autoregulation in 3-phosphoinositide-dependent Grp1 family Arf GTPase exchange factors. Mol Cell 28, 569–83 (2007).

10. He, X., Kuo, Y.C., Rosche, T.J. & Zhang, X. Structural basis for autoinhibition of the guanine nucleotide exchange factor FARP2. Structure 21, 355–64 (2013).

11. Rossman, K.L. et al. A crystallographic view of interactions between Dbs and Cdc42: PH domain-assisted guanine nucleotide exchange. EMBO J 21, 1315–26 (2002).

12. Jian, X.Y. et al. Molecular Basis for Cooperative Binding of Anionic Phospholipids to the PH Domain of the Arf GAP ASAP1. Structure 23, 1977–1988 (2015).

13. Kam, J.L. et al. Phosphoinositide-dependent activation of the ADP-ribosylation factor GTPase-activating protein ASAP1 - Evidence for the pleckstrin homology domain functioning as an allosteric site. Journal of Biological Chemistry 275, 9653–9663 (2000).

14. Roy, N.S. et al. Interaction of the N terminus of ADP-ribosylation factor with the PH domain of the GTPase-activating protein ASAP1 requires phosphatidylinositol 4,5-bisphosphate. J Biol Chem 294, 17354–17370 (2019).

15. Soubias, O. et al. Membrane surface recognition by the ASAP1 PH domain and consequences for interactions with the small GTPase Arf1. Sci Adv 6(2020).

16. Denisov, I.G. & Sligar, S.G. Nanodiscs in Membrane Biochemistry and Biophysics. Chem Rev 117, 4669–4713 (2017).

17. Hagn, F., Etzkorn, M., Raschle, T. & Wagner, G. Optimized phospholipid bilayer nanodiscs facilitate high-resolution structure determination of membrane proteins. J Am Chem Soc 135, 1919–25 (2013).

18. Antonny, B., Beraud-Dufour, S., Chardin, P. & Chabre, M. N-terminal hydrophobic residues of the G-protein ADP-ribosylation factor-1 insert into membrane phospholipids upon GDP to GTP exchange. Biochemistry 36, 4675–84 (1997).

19. Kahn, R.A. & Gilman, A.G. The protein cofactor necessary for ADP-ribosylation of Gs by cholera toxin is itself a GTP binding protein. J Biol Chem 261, 7906–11 (1986).

20. Li, Y. et al. Functional Expression and Characterization of Human Myristoylated-Arf1 in Nanodisc Membrane Mimetics. Biochemistry 58, 1423–1431 (2019).

21. Randazzo, P.A., Miura, K. & Jackson, T.R. Assay and purification of phosphoinositide-dependent ADP-ribosylation factor (ARF) GTPase activating proteins. *Regulators and Effectors of Small Gtpases*, Pt E 329, 343–354 (2001).

22. Luo, R.B. et al. Kinetic analysis of GTP hydrolysis catalysed by the Arf1-GTP-ASAP1 complex. Biochemical Journal 402, 439–447 (2007).

23. Pervushin, K., Riek, R., Wider, G. & Wuthrich, K. Attenuated T2 relaxation by mutual cancellation of dipole-dipole coupling and chemical shift anisotropy indicates an avenue to NMR structures of very large biological macromolecules in solution. Proc Natl Acad Sci U S A 94, 12366–71 (1997).

24. Hamel, D.J. & Dahlquist, F.W. The contact interface of a 120 kD CheA-CheW complex by methyl TROSY interaction spectroscopy. J Am Chem Soc 127, 9676–7 (2005).

25. Battiste, J.L. & Wagner, G. Utilization of site-directed spin labeling and high-resolution heteronuclear nuclear magnetic resonance for global fold determination of large proteins with limited nuclear overhauser effect data. Biochemistry 39, 5355–65 (2000).

26. Clore, G.M. Practical Aspects of Paramagnetic Relaxation Enhancement in Biological Macromolecules. Methods Enzymol 564, 485–97 (2015).

27. Dominguez, C., Boelens, R. & Bonvin, A.M. HADDOCK: a protein-protein docking approach based on biochemical or biophysical information. J Am Chem Soc 125, 1731–7 (2003).

28. van Zundert, G.C.P. et al. The HADDOCK2.2 Web Server: User-Friendly Integrative Modeling of Biomolecular Complexes. J Mol Biol 428, 720–725 (2016).

29. White, A.D. et al. Free energy of solvated salt bridges: a simulation and experimental study. J Phys Chem B 117, 7254–9 (2013).

30. Li, J. et al. Optimization of sortase A ligation for flexible engineering of complex protein systems. J Biol Chem 295, 2664–2675 (2020).

31. Zhang, Y. et al. Myr-Arf1 conformational flexibility at the membrane surface sheds light on the interactions with ArfGAP ASAP1. Nat Commun 14, 7570 (2023).

32. Calixto, A.R., Moreira, C. & Kamerlin, S.C.L. Recent Advances in Understanding Biological GTP Hydrolysis through Molecular Simulation. ACS Omega 5, 4380–4385 (2020).

33. Gerwert, K., Mann, D. & Kotting, C. Common mechanisms of catalysis in small and heterotrimeric GTPases and their respective GAPs. Biol Chem 398, 523–533 (2017).

34. Scheffzek, K. et al. The Ras-RasGAP complex: structural basis for GTPase activation and its loss in oncogenic Ras mutants. Science 277, 333–8 (1997).

35. Kremer, W., Steiner, G., Beraud-Dufour, S. & Kalbitzer, H.R. Conformational states of the small G protein Arf-1 in complex with the guanine nucleotide exchange factor ARNO-Sec7. J Biol Chem 279, 17004–12 (2004).

36. Mishra, B. & Johnson, M.E. Speed limits of protein assembly with reversible membrane localization. J Chem Phys 154, 194101 (2021).

37. Wu, Y., Vendome, J., Shapiro, L., Ben-Shaul, A. & Honig, B. Transforming binding affinities from three dimensions to two with application to cadherin clustering. Nature 475, 510–3 (2011).

38. Yogurtcu, O.N. & Johnson, M.E. Cytosolic proteins can exploit membrane localization to trigger functional assembly. PLoS Comput Biol 14, e1006031 (2018).

39. Siemers, M. et al. Bridge: A Graph-Based Algorithm to Analyze Dynamic H-Bond Networks in Membrane Proteins. J Chem Theory Comput 15, 6781–6798 (2019).

40. Venkatakrishnan, A.J. et al. Diverse GPCRs exhibit conserved water networks for stabilization and activation. Proc Natl Acad Sci U S A 116, 3288–3293 (2019).

41. Scheffzek, K. & Shivalingaiah, G. Ras-Specific GTPase-Activating Proteins-Structures, Mechanisms, and Interactions. Cold Spring Harb Perspect Med 9(2019).

42. Ahmadian, M.R., Stege, P., Scheffzek, K. & Wittinghofer, A. Confirmation of the arginine-finger hypothesis for the GAP-stimulated GTP-hydrolysis reaction of Ras. Nat Struct Biol 4, 686–9 (1997).

43. Ahmadian, M.R. et al. Guanosine triphosphatase stimulation of oncogenic Ras mutants. Proc Natl Acad Sci U S A 96, 7065–70 (1999).

44. Ismail, S.A., Vetter, I.R., Sot, B. & Wittinghofer, A. The structure of an Arf-ArfGAP complex reveals a Ca2+ regulatory mechanism. Cell 141, 812–21 (2010).

45. Menetrey, J. et al. Structural basis for ARF1-mediated recruitment of ARHGAP21 to Golgi membranes. EMBO J 26, 1953–62 (2007).

46. Malaby, A.W., van den Berg, B. & Lambright, D.G. Structural basis for membrane recruitment and allosteric activation of cytohesin family Arf GTPase exchange factors. Proc Natl Acad Sci U S A 110, 14213–8 (2013).

47. Liu, Y., Kahn, R.A. & Prestegard, J.H. Interaction of Fapp1 with Arf1 and PI4P at a membrane surface: an example of coincidence detection. Structure 22, 421–30 (2014).

48. Jiao, Y. et al. Genipin, a natural AKT inhibitor, targets the PH domain to affect downstream signaling and alleviates inflammation. Biochem Pharmacol 170, 113660 (2019).

49. Kang, Y. et al. Regulation of AKT Activity by Inhibition of the Pleckstrin Homology Domain-PtdIns(3,4,5)P(3) Interaction Using Flavonoids. J Microbiol Biotechnol 28, 1401–1411 (2018).

50. Kumar, C.C. & Madison, V. AKT crystal structure and AKT-specific inhibitors. Oncogene 24, 7493–501 (2005).

51. Mahadevan, D. et al. Discovery of a novel class of AKT pleckstrin homology domain inhibitors. Mol Cancer Ther 7, 2621–32 (2008).

52. Meuillet, E.J. Novel inhibitors of AKT: assessment of a different approach targeting the pleckstrin homology domain. Curr Med Chem 18, 2727–42 (2011).

53. Cash, J.N. et al. Discovery of Small Molecules That Target the Phosphatidylinositol (3,4,5) Trisphosphate (PIP(3))-Dependent Rac Exchanger 1 (P-Rex1) PIP(3)-Binding Site and Inhibit P-Rex1-Dependent Functions in Neutrophils. Mol Pharmacol 97, 226–236 (2020).

54. Nawrotek, A. et al. PH-domain-binding inhibitors of nucleotide exchange factor BRAG2 disrupt Arf GTPase signaling. Nat Chem Biol 15, 358–366 (2019).

55. Rosenberg, E.M., Jr. et al. The small molecule inhibitor NAV-2729 has a complex target profile including multiple ADP-ribosylation factor regulatory proteins. J Biol Chem 299, 102992 (2023).

56. Luo, R., Ha, V.L., Hayashi, R. & Randazzo, P.A. Arf GAP2 is positively regulated by coatomer and cargo. Cellular Signalling 21, 1169–1179 (2009).

57. Luo, R., Jenkins, L.M.M., Randazzo, P.A. & Gruschus, J. Dynamic interaction between Arf GAP and PH domains of ASAP1 in the regulation of GAP activity. Cellular Signalling 20, 1968–1977 (2008).

58. Pettersen, E.F. et al. UCSF Chimera--a visualization system for exploratory research and analysis. J Comput Chem 25, 1605–12 (2004).

59. Li, J. & Byrd, R.A. A simple protocol for the production of highly deuterated proteins for biophysical studies. J Biol Chem 298, 102253 (2022).

60. Delaglio, F. et al. NMRPipe: a multidimensional spectral processing system based on UNIX pipes. J Biomol NMR 6, 277–93 (1995).

61. Favier, A. & Brutscher, B. NMRlib: user-friendly pulse sequence tools for Bruker NMR spectrometers. J Biomol NMR 73, 199–211 (2019).

62. Jumper, J. et al. Highly accurate protein structure prediction with AlphaFold. Nature 596, 583–589 (2021).

63. Jo, S., Kim, T., Iyer, V.G. & Im, W. CHARMM-GUI: a web-based graphical user interface for CHARMM. J Comput Chem 29, 1859–65 (2008).

64. Huang, J. et al. CHARMM36m: an improved force field for folded and intrinsically disordered proteins. Nat Methods 14, 71–73 (2017).

65. Klauda, J.B. et al. Update of the CHARMM all-atom additive force field for lipids: validation on six lipid types. J Phys Chem B 114, 7830–43 (2010).

66. Michaud-Agrawal, N., Denning, E.J., Woolf, T.B. & Beckstein, O. Software News and Updates MDAnalysis: A Toolkit for the Analysis of Molecular Dynamics Simulations. Journal of Computational Chemistry 32, 2319–2327 (2011).

67. Polyhach, Y., Bordignon, E. & Jeschke, G. Rotamer libraries of spin labelled cysteines for protein studies. Physical Chemistry Chemical Physics 13, 2356–2366 (2011).

