## supplemental information for "The PH domain in the ArfGAP ASAP1 drives catalytic activation through an unprecedented allosteric mechanism"

FIG.SI1

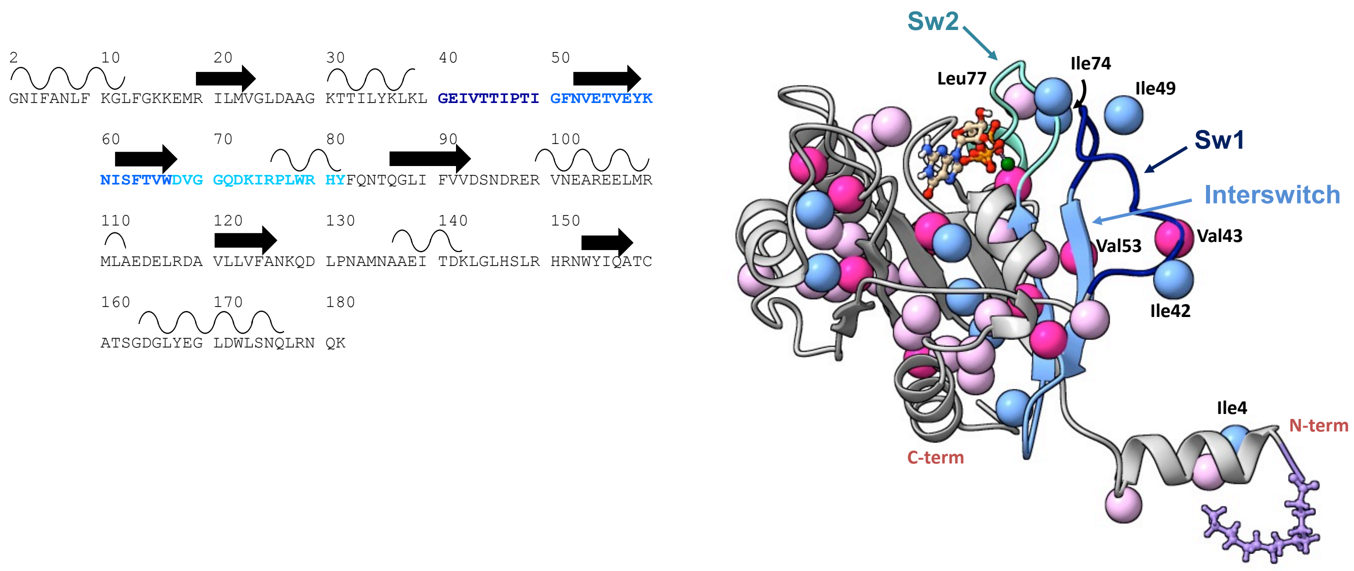


**Fig. SI1. A)** Amino acid sequence and secondary structure elements of Arf1. Switch 1 (dark blue), switch 2 (cyan) and the IS (corn blue) are highlighted **B)** Homology model of human myr-Arf1, generated based on yeast Arf1 (PDB:2KSQ) using MODELLER, is shown in grey ribbon format. Switch 1 (dark blue), switch 2 (cyan) and the interswitch (light blue) and methyl-containing residues are highlighted: 11 isoleucines (blue), 22 leucines (light pink), and 11 valines (dark pink). The myristoyl chain (purple) is shown as a ball and stick representation. Residues 2–13 form the N-terminal helix (embedded in the nanodisc), and residues 17–181 constitute the G-domain (solvent-exposed). For leucine and valine residues, only the Pro-S methyl carbons are shown. Images created using Chimera.

FIG.SI2

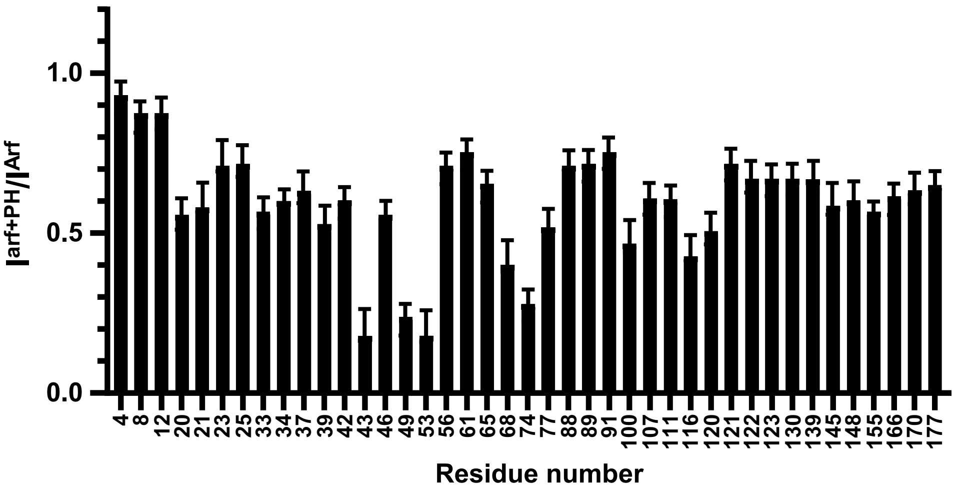


**Fig. SI2.** Attenuation of Arf methyl resonances upon complex formation. Ratio of cross peak intensity $I^{Arf+PH}$ / $I^{Arf}$ observed for methyl residues with ($I^{Arf+PH}$) and without ($I^{Arf}$) PH domain. When in complex with *wt* ASAP1 PH, loss of rotational freedom leads to an average resonance attenuation of 0.65 ± 0.07. Additional selective resonance attenuation observed for residues 43 and 49 of switch 1, 53 of the interswitch and 68 and 74 of switch 2 reflect residues at the interface.

FIG.SI3

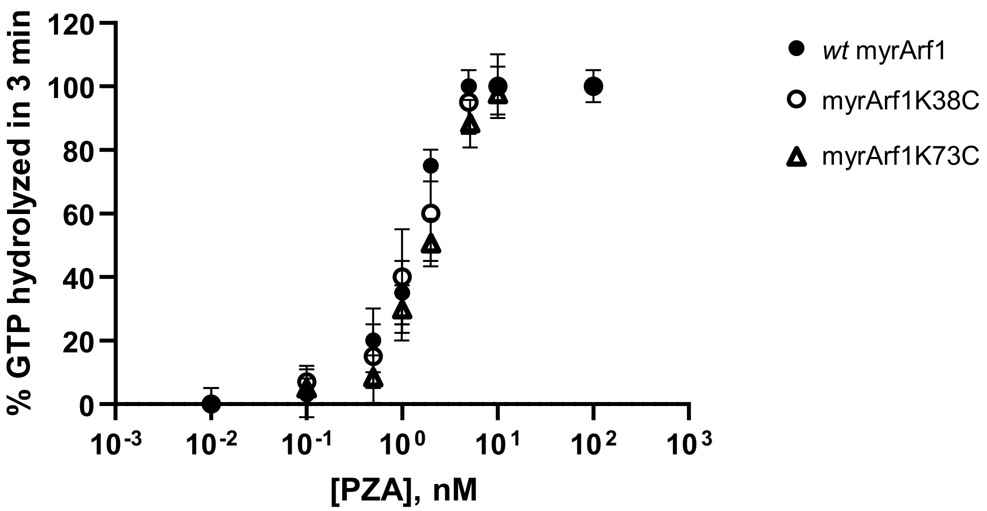


**Fig. SI3.** Functional activity of myrArf1, myrArf1K73C or myrArf1K38C. *wt* PZA was titrated into a reaction containing 5 μM myrArf1·GTP, myrArf1K73C·GTP or myrArf1K38C·GTP as a substrate. The percentage of GTP bound to myr-Arf1 hydrolyzed in 3 min is plotted against *wt* PZA concentration.

FIG.SI4

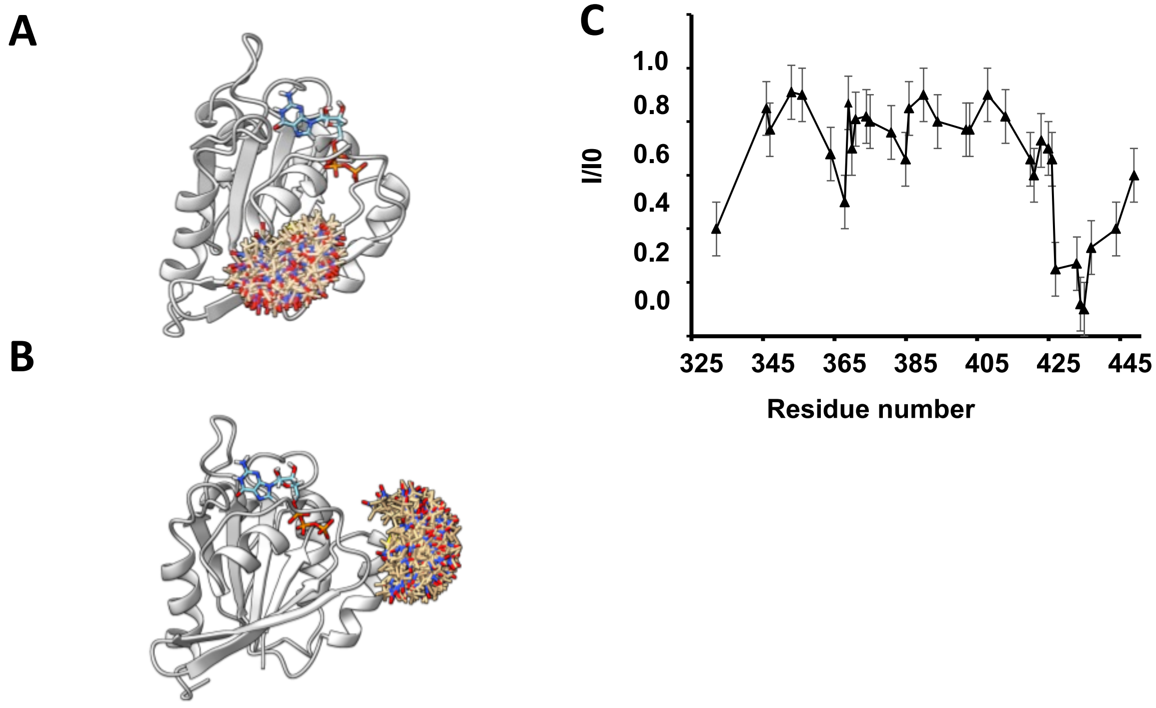


**Fig. SI4. A)** Position of MTSL label (brown) on myrArfK38C (in grey). For simplicity, residue 2-17 were omitted. Motion of the label is indicated by the multiple positions depicted. **B)** Position of MTSL label (brown) on myrArfK73C (in grey). For simplicity, Arf is represented without residue 2-17. **C)** Intermolecular PRE profile measured on ASAP1 PH in the presence of MTSL-tagged myrArf1K73C at the membrane surface. Two independent experiments were performed. Data are presented as mean values. Error bars were calculated based on the signal-to-noise (S/N) ratio of the spectra as described in Methods.

FIG.SI5

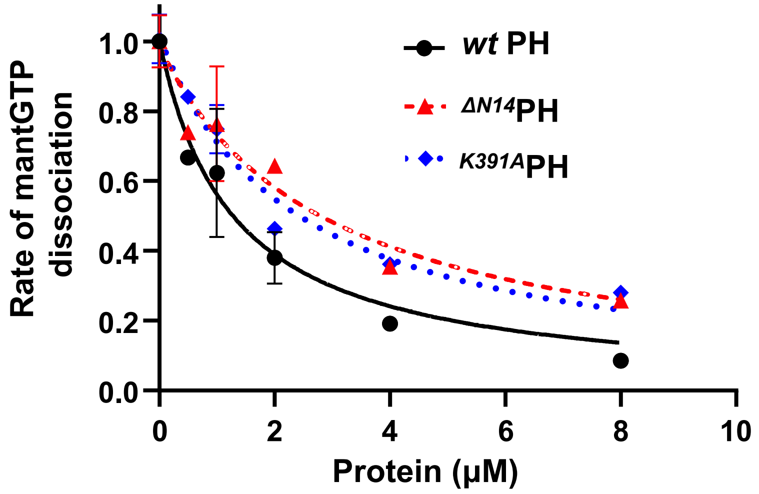


**Fig. SI5.** Dissociation rates of mantGTP as a function of PH domain concentration for *wt* PH (black circle), $^{\Delta N14}$PH (red triangle) and $^{K391A}$PH (blue diamond). The dissociation rates are normalized to the rate measured in the absence of PH domain (0.05 min$^{-1}$).

FIG.SI6

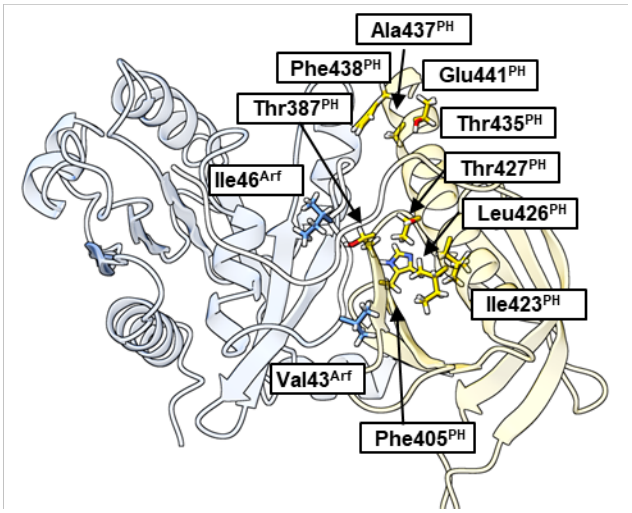


**Fig. SI6. A)** Position of additional mutated residues on myrArf (blue) and ASAP1 PH (gold). See also Tables SI3-5 and Fig. 5D.

FIG.SI7

**A**

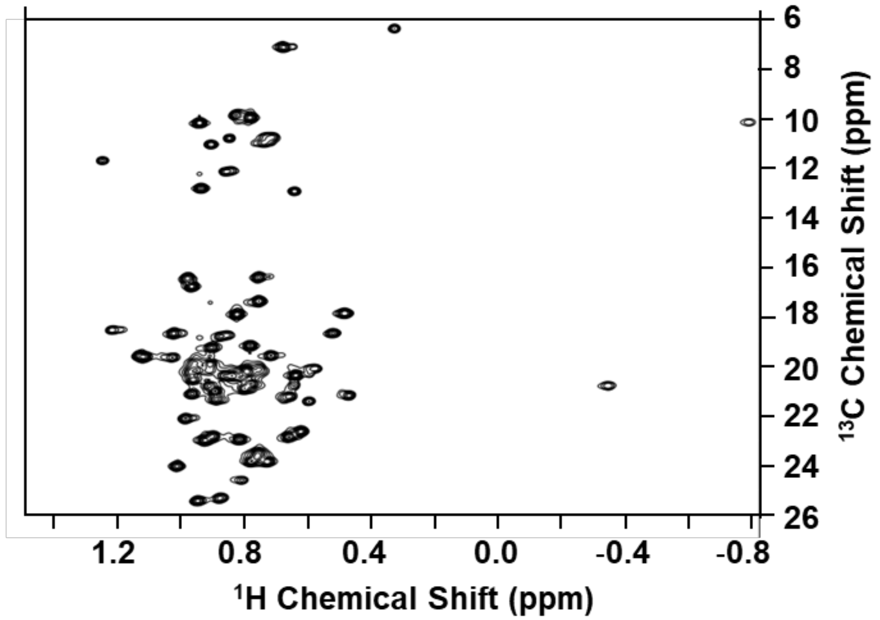


**B**

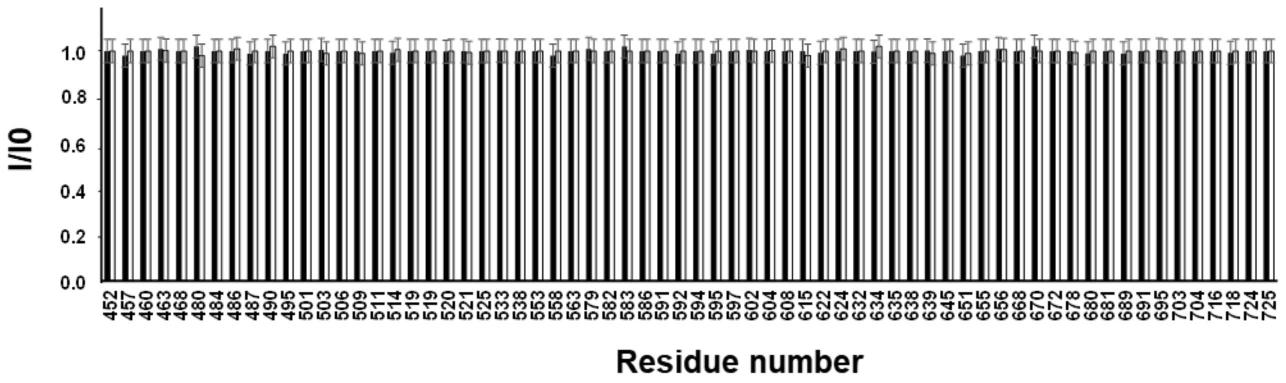


**Fig. SI7. A)** $^{1}$H-$^{13}$C HMQC of U-$^{2}$H,$^{15}$N and δ1-$^{13}$C$^{1}$H-labeled Ile, δ1 -$^{13}$C$^{1}$H-labeled Leu and γ1-$^{13}$C$^{1}$H-labeled Val ZA domain (100 μM) in the presence of MTSL-tagged myrArf1C159

(100μM) bound to equimolar ratio of ASAP1 PH at the membrane surface. **B)** Intermolecular PRE ratio measured on ZA in the presence of MTSL-tagged myrArf1C159 alone (black column) or bound to equimolar ratio of ASAP1 PH (open column) at the membrane surface. No significant PRE could be detected.

FIG.SI8

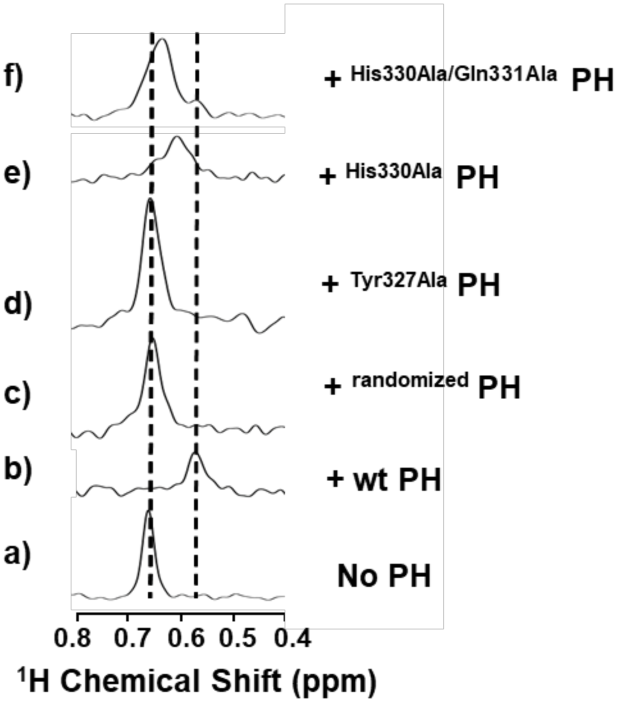


**Fig. SI8.** Stack of rows extracted from a $^{1}H$-$^{13}C$ HMQC experiment along the proton dimension of Val43 (myrArf1) in the absence (a) or in the presence of wt-ASAP1 PH (b), ASAP1 PH with a randomized N-term extension (c) Tyr327Ala ASAP1 PH (d) His330Ala ASAP1 PH (e) or His330Ala/Gln331Ala ASAP1 PH (f).

### TABLE SI1

| | C50 (in M) | |
|---|---|---|
| **Protein** | **+ PI(4,5)$P_2$** | **- PI(4,5)$P_2$** |
| *wt* PZA | $3.10^{-11} \pm 1.5.10^{-11}$ (7) | $1.10^{-6} \pm 1.6.10^{-6}$ (3) |
| $^{\Delta N14}$ PZA | $3.10^{-9} \pm 1.8.10^{-9}$ (3) | ND |
| PHdZA | $3.10^{-7} \pm 1.7.10^{-7}$ (3) | $>1.10^{-3}$ (3) |
| ZA | $> 0.12.10^{-3}$ (4) | $> 0.12.10^{-3}$ (3) |
| *wt* PH + ZA | $4.1.10^{-6} \pm 0.3.10^{-6}$ (4) | ND |
| $^{\Delta N14}$PH + ZA | $2.10^{-4} \pm 1.7.10^{-4}$ (4) | ND |

**Table SI1.** Comparison of GAP activity using Arf1 as substrate in membranes with or without PI(4,5)$P_2$. The amount of GAP (in M) required to achieve 50% conversion of [$\alpha^{32}$P]GTP to [$\alpha^{32}$P]GDP in 3 min ($C_{50}$) was estimated and is inversely proportional to enzymatic power. Data are expressed as mean values ± SD with the number of repeats indicated between parentheses.

### TABLE SI2

**Ambiguous interaction restraints used in the structure calculations of Clusters$^{\beta}{}_{5}{}^{/\beta}{}_{7}$**

| Protein | Active Residues | Passive Residues |
|---|---|---|
| myrArf1 | 20, 40, 47, 49, 52, 53, 77, 79, 84 | 51, 55 |
| ASAP1 PH | 387, 388, 389, 390, 408, 423, 434, 435, 437 | none |

**Ambiguous interaction restraints used in the structure calculations of Clusters$^{\beta}{}_{2}{}^{/\beta}{}_{3}$**

| Protein | Active Residues | Passive Residues |
|---|---|---|
| myrArf1 | 20, 40, 47, 49, 52, 53, 77, 79, 84 | 51, 55 |
| ASAP1 PH | 368, 370 | 364, 371 |

**Unambiguous Distance restraints from PRE measurements used in the structure calculation of Clusters$^{\beta}{}_{5}{}^{/\beta}{}_{7}$**

| Residue | Sγ atom | Distance (Å) |
|---|---|---|
| Ile403 | K38C | 1.8 - 16 |
| Thr408 | K38C | 1.8 - 16 |
| Ala394 | K38C | 1.8 - 16 |

**Unambiguous Distance restraints from PRE measurements used in the structure calculation of Clusters$^{\beta}{}_{2}{}^{/\beta}{}_{3}$**

| Residue | Sγ atom | Distance (Å) |
|---|---|---|
| Ala363 | K38C | 1.8 - 16 |
| Ile368 | K38C | 1.8 - 16 |
| Thr370 | K38C | 1.8 - 16 |

### TABLE SI3

| myrArf1 protein | GAP assays | |
|---|---|---|
| | $C_{50}$ (nM) | Fold change |
| **WT** | 0.036 ± 0.014 (7) | 1 |
| **E41A** | 0.11 ± 0.02 (3) | 3.2 |
| **I46A** | 48 ± 2 (3) | 1337 |
| **V43A** | 0.25 ± 0.04 (4) | 6.9 |
| **E54A** | 2.57 ± 0.32 (3) | 72 |

**Table SI3.** Comparison of GAP activity for myrArf1 mutants measured in LUV containing 5 mol% of PI(4,5)$P_2$. The amount of GAP (in nM) required to achieve 50% conversion of [$\alpha^{32}$P]GTP to [$\alpha^{32}$P]GDP in 3 min ($C_{50}$) was estimated. Data are expressed as mean values ± SD with the number of repeats indicated between parentheses.

### TABLE SI4

| ASAP1 PZA | GAP assays | |
|---|---|---|
| | $C_{50}$ (nM) | Fold change |
| **WT** | 0.036 ± 0.014 (7) | 1 |
| **T387L** | 0.021 ± 0.005 (4) | 0.6 |
| **K391A** | 3.58 ± 1.055 (4) | 99.8 |
| **H405E** | 0.154 ± 0.069 (6) | 4.3 |
| **F438A** | 0.564 ± 0.28 (4) | 15.7 |
| **E441A** | 0.023± 0.005 (4) | 1.2 |
| **E441R** | 0.045± 0.008 (4) | 0.6 |

**Table SI4.** Comparison of GAP activity for PZA mutants measured in LUV containing 5 mol% of $PI(4,5)P_2$. The amount of GAP (in nM) required to achieve 50% conversion of [$\alpha^{32}$P]GTP to [$\alpha^{32}$P]GDP in 3 min ($C_{50}$) was estimated. Data are expressed as mean values ± SD with the number of repeats indicated between parentheses.

### TABLE SI5

| ASAP1 PZA | GAP assays | |
|---|---|---|
| | $C_{50}$ (nM) | Fold change |
| **WT*** | 0.36 ± 0.14 (7) | 1 |
| **I423A*** | 0.756 ± 0.17 (3) | 2.1 |
| **L426A*** | 1.86 ± 0.23 (3) | 5.19 |
| **T427A*** | 0.154 ± 0.1 (3) | 0.43 |
| **T435A*** | 0.129 ± 0.1 (3) | 0.36 |
| **A437R*** | 1.303± 0.1 (3) | 3.62 |

**Table SI5.** Comparison of GAP activity for PZA mutants measured in LUVs at 1 mol% PI(4,5)$P^2$. The amount of GAP (in nM) required to achieve 50% conversion of [$\alpha^{32}$P]GTP to [$\alpha^{32}$P]GDP in 3 min ($C_{50}$) was estimated. Data are expressed as mean values ± SD with the number of repeats indicated between parentheses. Asterisk indicated determination was made with 1 mol% PIP2 in the LUVs. All other experiments with LUVs included 5 mol% PIP2.

### KINETIC MODELING

"In Trans" Reaction Network

Our model reaction network for the in trans experiments tracks the concentrations of 11 species in total. The 4 fundamental species are PH, ZA, ArfGTP, and $PIP_2$. Species formed through bimolecular association reactions are PH•ArfGTP, ArfGTP•ZA, PH•ArfGTP•ZA, $PH_m$ (PH which has bound $PIP_2$ and is on the membrane), $PH_m$•ArfGTP, and $PH_m$•ArfGTP•ZA. Finally, there is the product ArfGDP formed through GTP hydrolysis. The complete list of reactions, along with associated rate constants, is:

1. $\mathrm{ArfGTP} + \mathrm{ZA} \rightleftharpoons \mathrm{ArfGTP} \bullet \mathrm{ZA} \ : \ k_{on/off}^{\mathrm{Arf+ZA}}$
2. $\mathrm{ArfGTP} + \mathrm{PH} \rightleftharpoons \mathrm{PH} \bullet \mathrm{ArfGTP} \ : \ k_{on/off}^{\mathrm{Arf+PH}}$
3. $\mathrm{PH} \bullet \mathrm{ArfGTP} + \mathrm{ZA} \rightleftharpoons \mathrm{PH} \bullet \mathrm{ArfGTP} \bullet \mathrm{ZA} \ : \ k_{on/off}^{\mathrm{ZA}}$
4. $\mathrm{PIP_2} + \mathrm{PH} \rightleftharpoons \mathrm{PH_m} \ : \ k_{on/off}^{\mathrm{PH+PIP}}$
5. $\mathrm{PH_m} \bullet \mathrm{ArfGTP} + \mathrm{ZA} \rightleftharpoons \mathrm{PH_m} \bullet \mathrm{ArfGTP} \bullet \mathrm{ZA} \ : \ k_{on/off}^{\mathrm{ZA}}$
6. $\mathrm{ArfGTP} \bullet \mathrm{ZA} + \mathrm{PH} \rightleftharpoons \mathrm{PH} \bullet \mathrm{ArfGTP} \bullet \mathrm{ZA} \ : \ k_{on/off}^{\mathrm{Arf+PH}}$
7. $\mathrm{ArfGTP} + \mathrm{PH_m} \rightleftharpoons \mathrm{PH_m}.\mathrm{ArfGTP} \ : \ k_{on/off}^{\mathrm{Arf+PH}}$ (2D)
8. $\mathrm{ArfGTP} \bullet \mathrm{ZA} + \mathrm{PH_m} \rightleftharpoons \mathrm{PH_m} \bullet \mathrm{ArfGTP} \bullet \mathrm{ZA} \ : \ k_{on/off}^{\mathrm{Arf+PH}}$ (2D)
9. $\mathrm{PH} \bullet \mathrm{ArfGTP} + \mathrm{PIP_2} \rightleftharpoons \mathrm{PH_m} \bullet \mathrm{ArfGTP} \ : \ k_{on/off}^{\mathrm{PH+PIP}}$ (2D)
10. $\mathrm{PH} \bullet \mathrm{ArfGTP} \bullet \mathrm{ZA} + \mathrm{PIP_2} \rightleftharpoons \mathrm{PH_m} \bullet \mathrm{ArfGTP} \bullet ZA \ : \ k_{on/off}^{\mathrm{PH+PIP}}$ (2D)
11. $\mathrm{ArfGTP} \bullet \mathrm{ZA} \rightarrow \mathrm{ArfGDP} + \mathrm{ZA} \ : \ k_{cat}^{\mathrm{Arf.ZA}}$
12. $\mathrm{PH} \bullet \mathrm{ArfGTP} \bullet \mathrm{ZA} \rightarrow \mathrm{ArfGDP} + \mathrm{PH} + \mathrm{ZA} \ : \ k_{cat}^{\mathrm{PH.Arf.ZA}}$

13. $\mathrm{PH_m} \bullet \mathrm{ArfGTP} \bullet \mathrm{ZA} \rightarrow \mathrm{ArfGDP} + \mathrm{PH_m} + \mathrm{ZA}$ : $k_{cat}^{\mathrm{PHm.Arf.ZA}}$

The associated system of ODEs is given at the end of this document.

ArfGTP is assumed to reside entirely on the membrane due to its N-terminal myristoylation. Rates followed by "(2D)" indicate reactions which occur on the two-dimensional membrane surface. The intrinsic (microscopic) association rates for 2D reactions (dimensions $area^{-1}time^{-1}$) are related to their 3D counterparts (dimensions $volume^{-1}time^{-1}$) by dividing by a nanoscopic length scale $h$ which phenomenologically accounts for thermodynamic effects of surface binding. $h$ is typically comparable to the molecular length scale, i.e., of order nanometers [5]. Since in our ODE system we track all species based on volume concentration, the effective macroscopic association rates (dimensions $volume^{-1}time^{-1}$) for 2D reactions used in these equations are multiplied by a unitless *dimensionality factor* $\mathrm{DF} = V/Ah$, where $V$ is the solution volume and $A$ is the total membrane area. Critically, the $k_{cat}$ rate is allowed to change based on the presence of PH in complex with Arf and its $PIP_2$ binding state. Some simplifying assumptions made in the modeling are that the rates of association/dissociation for PH to Arf are independent of whether or not $PIP_2$ is bound, and that the catalytic rate $k_{cat}^{\mathrm{PHm.Arf.ZA}}$ is equal to the previously measured experimental value of approximately 56 $s^{-1}$. Additionally, the association of PH to the membrane via the binding of $PIP_2$ is modeled as a single bimolecular association reaction.

Tandem network

The tandem PZA reaction network is qualitatively different from the "in trans" system due to the presence of the linker between the PH and ZA subunits. After the binding of either the PH or ZA subunit to Arf, the subsequent binding of the other subunit becomes a first-order reaction rather than second-order. We introduce rates for these “loop-closure” reactions in the following way:

The dissociation constants for the individual subunits binding to Arf can be expressed as

$$K_d^{\text{Arf+ZA}} = \frac{k_{off}^{\text{Arf+ZA}}}{k_{on}^{\text{Arf+ZA}}} = c_0 e^{\,\Delta G_{\text{ZA}}\,/k_{\text{B}}T} \qquad K_d^{\text{Arf+PH}} = \frac{k_{off}^{\text{Arf+PH}}}{k_{on}^{\text{Arf+PH}}} = c_0 e^{\,\Delta G_{\text{PH}}\,/k_{\text{B}}T}$$

where the $k_{on/off}$ rates are the same as those in the reactions above, $\Delta G_{\text{ZA/PH}}$ is the change in Gibbs free energy upon ZA/PH binding, and $c_0$ is the standard state concentration 1 M. The affinity of PZA for Arf can be expressed as

$$K_d^{\text{Arf+PZA}} = c_0 e^{\,(\Delta G_{\text{ZA}} + \Delta G_{\text{PH}} + \Delta G_{\text{coop}})/k_{\text{B}}T}$$

where $\Delta G_{\text{coop}}$ is a cooperative contribution to the free energy of binding for PH and ZA to Arf due to the linker connecting them. Substituting the expressions above for $\Delta G_{\text{ZA/PH}}$ in terms of rates, we arrive at

$$K_d^{\text{Arf+PZA}} = \frac{k_{off}^{\text{Arf+ZA}}\, k_{off}^{\text{Arf+PH}}}{c_0 k_{on}^{\text{Arf+ZA}}\, k_{on}^{\text{Arf+PH}}} e^{\Delta G_{\text{coop}}/k_{\text{B}}T}.$$

Considering the case where PH binds first and rearranging this expression we find,

$$K_d^{\text{Arf+PZA}} = \frac{k_{off}^{\text{Arf+PH}}}{k_{on}^{\text{Arf+PH}}} \frac{k_{off}^{\text{Arf+ZA}}}{c_0 k_{on}^{\text{Arf+ZA}}\, e^{-\Delta G_{\text{coop}}/k_{\text{B}}T}}.$$

So, assuming that the off-rate $k_{off}^{\text{Arf+ZA}}$ is unchanged, we can straightforwardly read off the first-order binding rate for ZA to “close the loop” to Arf after PH is already bound: $k_{close}^{\text{Arf+ZA}} = c_0 k_{on}^{\text{Arf+ZA}}\, e^{-\Delta G_{\text{coop}}/k_{\text{B}}T}$. Rearranging the terms produces the analogous expression for PH loop-closure. The reactions which use these rates are highlighted with an asterisk (*) in the list below. That the off-rate remains the same is a reasonable simplifying assumption that we make here;

however, this can be relaxed at the cost of introducing a new parameter determining what fraction of $c_0\, e^{-\Delta G_{\mathrm{coop}}/k_{\mathrm{B}}T}$ is attributed to the on vs. off rate. The complete list of reactions for the tandem system is then:

1. $\mathrm{PZA} + \mathrm{PIP_2} \rightleftharpoons \mathrm{PZA_m} \ :\ k_{on/off}^{\mathrm{PH+PIP}}$
2. $\mathrm{ArfGTP} + \mathrm{PZA} \rightleftharpoons \mathrm{PH} \bullet \mathrm{ArfGTP} \ :\ k_{on/off}^{\mathrm{Arf+PH}}$
3. $\mathrm{ArfGTP} + \mathrm{PZA} \rightleftharpoons \mathrm{ArfGTP} \bullet \mathrm{ZA} \ :\ k_{on/off}^{\mathrm{Arf+ZA}}$
4. $\mathrm{ArfGTP} + \mathrm{PZA_m} \rightleftharpoons \mathrm{PH_m}.\mathrm{ArfGTP} \ :\ k_{on/off}^{\mathrm{Arf+PH}}$ (2D)
5. $\mathrm{ArfGTP} + \mathrm{PZA_m} \rightleftharpoons \mathrm{ArfGTP} \bullet \mathrm{ZA} \ :\ k_{on/off}^{\mathrm{Arf+ZA}}$ (2D)
6. $\mathrm{PH} \bullet \mathrm{ArfGTP} + \mathrm{PIP_2} \rightleftharpoons \mathrm{PH_m} \bullet \mathrm{ArfGTP} \ :\ k_{on/off}^{\mathrm{PH+PIP}}$ (2D)
7. $\mathrm{PH} \bullet \mathrm{ArfGTP} \bullet \mathrm{ZA} + \mathrm{PIP_2} \rightleftharpoons \mathrm{PH_m} \bullet \mathrm{ArfGTP} \bullet \mathrm{ZA} \ :\ k_{on/off}^{\mathrm{PH+PIP}}$ (2D)
8. $\mathrm{ArfGTP} \bullet \mathrm{ZA} + \mathrm{PIP_2} \rightleftharpoons \mathrm{ArfGTP} \bullet \mathrm{ZA_m} \ :\ k_{on/off}^{\mathrm{PH+PIP}}$ (2D)
9. $\mathrm{PH} \bullet \mathrm{ArfGTP} \rightleftharpoons \mathrm{PH} \bullet \mathrm{ArfGTP} \bullet \mathrm{ZA} \ :\ k_{close/off}^{\mathrm{Arf+ZA}}$ *
10. $\mathrm{PH_m} \bullet \mathrm{ArfGTP} \rightleftharpoons \mathrm{PH_m} \bullet \mathrm{ArfGTP} \bullet \mathrm{ZA} \ :\ k_{close/off}^{\mathrm{Arf+ZA}}$ *
11. $\mathrm{ArfGTP} \bullet \mathrm{ZA} \rightleftharpoons \mathrm{PH} \bullet \mathrm{ArfGTP} \bullet \mathrm{ZA} \ :\ k_{close/off}^{\mathrm{Arf+PH}}$ *
12. $\mathrm{ArfGTP} \bullet \mathrm{ZA_m} \rightleftharpoons \mathrm{PH_m} \bullet \mathrm{ArfGTP} \bullet \mathrm{ZA} \ :\ k_{close/off}^{\mathrm{Arf+PH}}$*
13. $\mathrm{ArfGTP} \bullet \mathrm{ZA} \rightarrow \mathrm{ArfGDP} + \mathrm{PZA} \ :\ k_{cat}^{\mathrm{Arf.ZA}}$
14. $\mathrm{ArfGTP} \bullet \mathrm{ZA_m} \rightarrow \mathrm{ArfGDP} + \mathrm{PZA_m} \ :\ k_{cat}^{\mathrm{Arf.ZA}}$
15. $\mathrm{PH} \bullet \mathrm{ArfGTP} \bullet \mathrm{ZA} \rightarrow \mathrm{ArfGDP} + \mathrm{PZA} \ :\ k_{cat}^{\mathrm{PH.Arf.ZA}}$
16. $\mathrm{PH_m} \bullet \mathrm{ArfGTP} \bullet \mathrm{ZA} \rightarrow \mathrm{ArfGDP} + \mathrm{PZA_m} \ :\ k_{cat}^{\mathrm{PHm.Arf.ZA}}$

In the above list, the notation $\mathrm{ZA_m}$ is used when the ZA subunit is bound to Arf and PH is bound to $\mathrm{PIP_2}$, but the PH subunit is not bound to Arf. Once again, the associated system of ODEs is given at the end of this document.

Two further assumptions have been made such that the reaction rates for this system are determined entirely in terms of the rates for the in trans reactions and the new parameter $\Delta G_{\mathrm{coop}}$. First, as $PIP_2$ binding occurs on the PH domain, the rates for membrane association of PZA are taken equal to those for PH alone. Second, the $k_{cat}$ values are taken equal to those for the in trans system with the same subunits present.

The simplified PHdZA model (green curves in Fig. 8A and E) uses the same reaction network as PZA with modified kinetic parameters: PHd binding to Arf is disallowed, as PHd has negligible affinity for Arf, and therefore $k_{cat}$ is assumed to be always equal to $k_{cat}^{\mathrm{Arf.ZA}}$. The hypothetical PZA model in which the only function of $PIP_2$ is localization (dashed/purple curves in Fig. 8A) is also modeled using the same equations with altered kinetics. In that case, $k_{cat}^{\mathrm{PHm.Arf.ZA}}$ is set equal to $k_{cat}^{\mathrm{PH.Arf.ZA}}$ instead of the previously determined experimental value for $k_{cat}$.

*Parameter Optimization based on Experimental Data*

The model parameters are constrained by experimental GTP hydrolysis rates as determined in several different conditions for both in trans and tandem PZA, as well as PHdZA. For the in trans systems, these include data for fraction of GTP hydrolyzed in 3 minutes with a fixed concentration of PH and varying amount of ZA (Fig. 8B), fixed concentration of ZA and varying PH (Fig. 8C), and ZA alone with no PH present (Fig. 8D). For tandem PZA, we use fraction of GTP hydrolyzed in the absence of $PIP_2$ and with 5 μM $PIP_2$ present (Fig. 8E). Fig. 8E also shows the PHdZA data used for fitting (red), for which the conditions are the same as PZA with $PIP_2$ present.

Our model parameters were optimized by minimizing the $\chi^2$ residual between the simulation results and the observational data points, while imposing certain restrictions on the model parameters. In addition to the parameter limits given in Table SI6, the following constraints were imposed on dissociation constants $K_\mathrm{d} = k_\mathrm{off}/k_\mathrm{on}$ based on experimental measurements: $\mathrm{DF} \times K_\mathrm{d}^{\mathrm{PH+Arf}} \geq 1\mu\mathrm{M}$, $K_\mathrm{d}^{\mathrm{ZA+Arf}} \geq 100\mu\mathrm{M}$, and $K_\mathrm{d}^{\mathrm{ZA+Arf}} > K_\mathrm{d}^{\mathrm{PH+Arf}}$. Additionally, the following constraints were imposed on rate constants to avoid pathological results: $k_{cat}^{\mathrm{PH.Arf.ZA}} \geq k_{cat}^{\mathrm{Arf.ZA}}$ and $k_{on}^{\mathrm{PHm.Arf+ZA}} \geq k_{on}^{\mathrm{PH.Arf+ZA}}$. These restrictions, along with parameters which are held constant during fitting, are summarized in Table SI7.

In order to find optimal parameter combinations in the high-dimensional space, we performed stochastic global optimization via a genetic algorithm implemented in the Julia programming language using packages from the SciML ecosystem [1–4]. An initial population of 3000 sets of parameters (individual candidates in the evolutionary algorithm) is sampled uniformly in log-space from the allowable parameter ranges (Table SI6) and respecting the imposed constraints (Table SI7). The genetic algorithm then proceeds by iterating the following steps for 10 generations:

1. Generate an offspring population via crossover and mutation:
    a. Adjacent candidate pairs have 50% probability of swapping parameters via 2-point crossover.
    b. Each individual parameter within a candidate then has a 75% chance of being uniformly scaled by up to 50% in either direction (within the prescribed bounds).

   c. Additionally, the 5 best individuals from the previous generation continue on unmodified.

2. Evaluate each candidate's fitness as the $\chi^2$ residual between the numerical ODE result and the data in Fig. 8B-E.
3. Select the next generation by tournament: 10% of the population is selected at random, and the individual with highest fitness is added to the next generation. This is repeated until the next generation has equal size.

We repeated this process 12 times, yielding a pool of 36,000 candidate parameter values. The overall best 100 fits (excluding duplicates) are the curves plotted in Fig. 8 and their parameter distributions are presented in Fig. SI9. The parameter values for the single best fit are given in Table SI6.

The large sample of initial candidate points followed by subsequent generations of stochastic updates allows the algorithm to sample many regions of the parameter space in order to avoid becoming trapped in an initial local minimum of the fitness landscape. However, the stochastic nature of the algorithm, combined with the rough nature of the high-dimensional fitness landscape, does mean that repeated optimization runs generally do not converge to identical sets of optimal parameters. Applying a deterministic minimization algorithm (such as downhill simplex) to the optimal candidate post-GA does not resolve this issue. However, post-GA parameter values tend to cluster within particular ranges, as shown in Fig. SI9, especially for certain parameters to which the model is particularly sensitive, in particular the catalytic rates. While the model is apparently not directly sensitive to the rates $k_{on/off}^{\mathrm{Arf+PH}}$ and $k_{on/off}^{\mathrm{Arf+ZA}}$ beyond a certain threshold, their ratios (affinities) are much more well-defined, varying over less than an order of magnitude, as shown in Fig. SI10.

| Variable Fit Parameters | | | |
|---|---|---|---|
| **Parameter** | **Units** | **Allowed Range** | **Best Fit** |
| $k_{on}^{\mathrm{Arf+ZA}}$ | $\mu M^{-1}s^{-1}$ | $10^{-8}$ — 10 | 0.029 |
| $k_{off}^{\mathrm{Arf+ZA}}$ | $s^{-1}$ | $10^{-3}$ — $10^{5}$ | 600 |
| $k_{on}^{\mathrm{Arf+PH}}$ | $\mu M^{-1}s^{-1}$ | $10^{-8}$ — 10 | 5.6 |
| $k_{off}^{\mathrm{Arf+PH}}$ | $s^{-1}$ | $10^{-3}$ — $10^{5}$ | $3.6\times10^{4}$ |
| $k_{cat}^{\mathrm{Arf.ZA}}$ | $s^{-1}$ | $10^{-6}$ — 56 | 0.045 |
| $k_{cat}^{\mathrm{PH.Arf.ZA}}$ | $s^{-1}$ | $10^{-6}$ — 56 | 0.050 |
| $k_{on}^{\mathrm{PH+PIP}}$ | $\mu M^{-1}s^{-1}$ | $10^{-8}$ — 10 | 0.022 |
| $K_{d}^{\mathrm{PH+PIP}}$ | μM | 0.1 — 10 | 0.68 |
| $h$ | nm | 1 — 100 | 16 |
| $\exp(-\Delta G_{\mathrm{coop}} / k_{\mathrm{B}}T)$ | | $10^{-5}$ — $10^{5}$ | 13 |

**Table SI6.** Variable model parameters with allowable ranges and optimal values.

| Fixed Parameters / Constraints | |
|---|---|
| **Parameter** | **Value / Constraint** |
| $k_{cat}^{\mathrm{PHm.Arf.ZA}}$ | 56 $s^{-1}$ [7] |
| $\mathrm{DF} \times K_{\mathrm{d}}^{\mathrm{Arf+PH}}$ | ≥ 1 μM |
| $K_{\mathrm{d}}^{\mathrm{Arf+ZA}}$ | ≥ 100 μM; ≥ $K_{\mathrm{d}}^{\mathrm{Arf+PH}}$ |
| $k_{cat}^{\mathrm{PH.Arf.ZA}}$ | ≥ $k_{cat}^{\mathrm{Arf.ZA}}$ |
| $k_{on}^{\mathrm{PHm.Arf+ZA}}$ | ≥ $k_{on}^{\mathrm{PH.Arf+ZA}}$ |

**Table SI7.** Kinetic model parameters which are fixed or constrained.

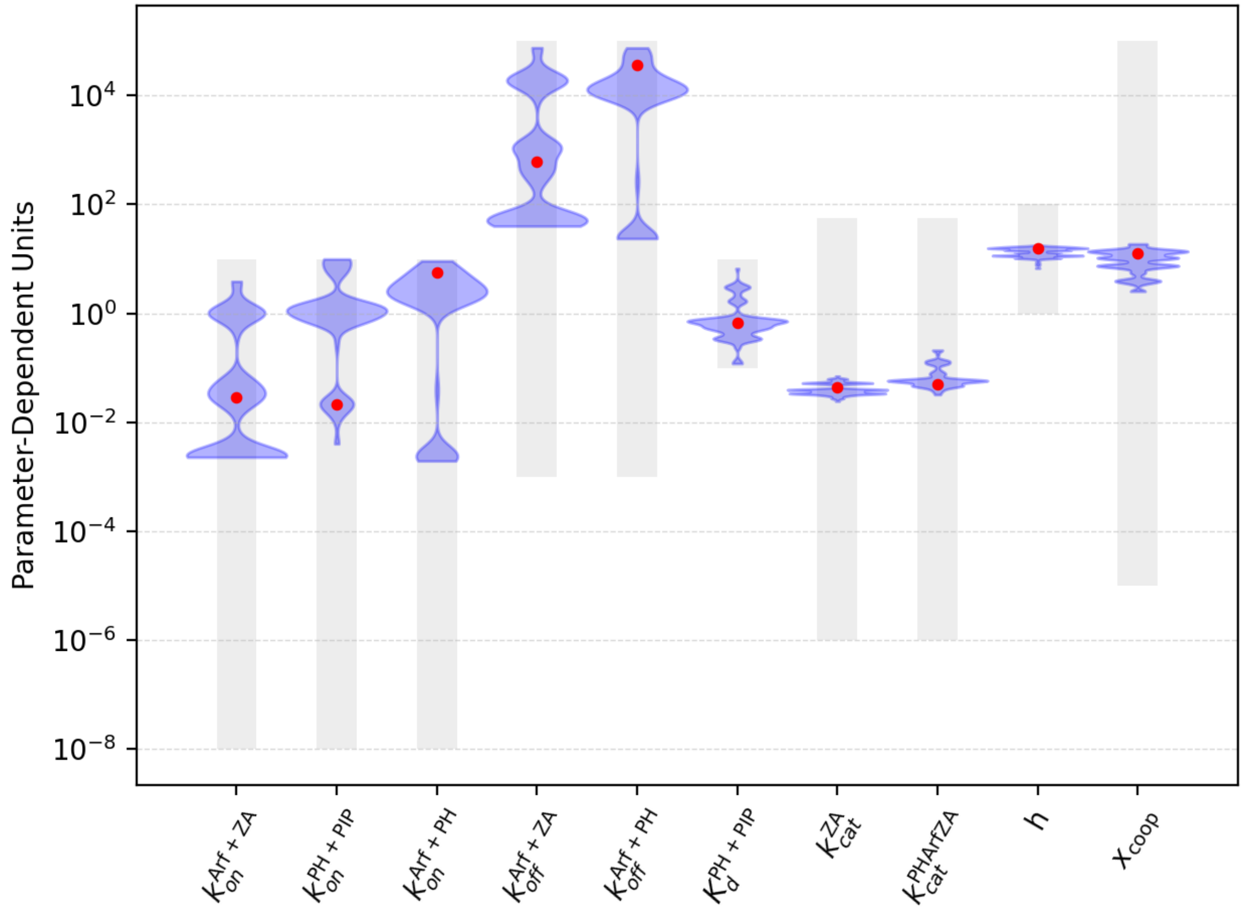


**Figure SI9.** Distribution of parameter values for 100 best solutions out of 36,000 from 12 runs of the GA. Units for each parameter are given in column 2 of Table SI6. The overall optimal parameter set (lowest $\chi^2$, values shown in Table SI6) is plotted as the red points. Grey shaded regions indicate allowed parameter ranges during optimization (column 3 of Table SI6). Parameter $x_{\mathrm{coop}} = \exp(-\Delta G_{\mathrm{coop}} \,/\, k_{\mathrm{B}}T\,)$.

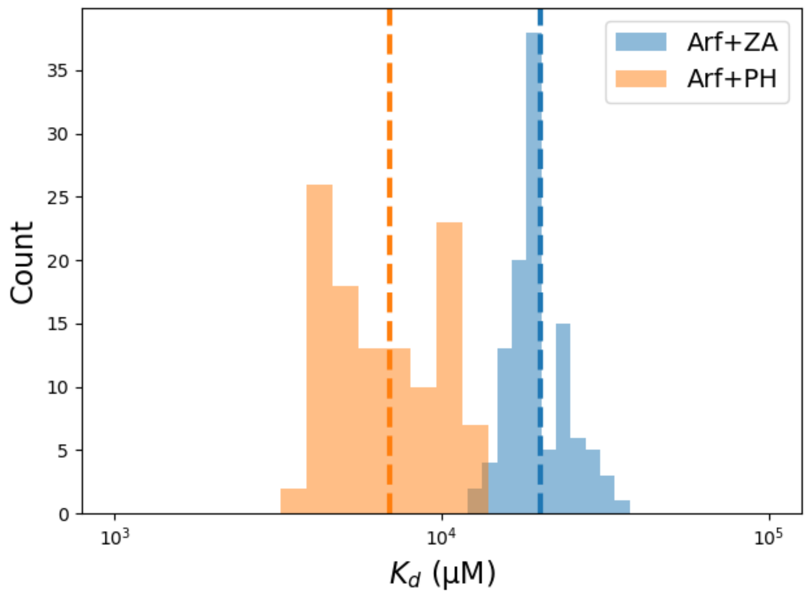


**Figure SI10.** Distribution of values for $K_\mathrm{d}^{\mathrm{Arf+PH}}$ and $K_\mathrm{d}^{\mathrm{Arf+ZA}}$ for the 100 best solutions (same as Fig. SI9). Dashed vertical lines indicate the overall best fit values (corresponding to the red points in Fig. SI9 and values given in Table SI6).

### 1 PH+ZA "in trans" Reaction ODEs

$$\frac{\mathrm{d[PIP_2]}}{\mathrm{d}t} = k_{off}^{\mathrm{PH+PIP}}[\mathrm{PH_m.Arf.ZA}] + k_{off}^{\mathrm{PH+PIP}}[\mathrm{PH_m}] + k_{off}^{\mathrm{PH+PIP}}[\mathrm{PH_m.Arf}] - k_{on}^{\mathrm{PH+PIP}}[\mathrm{PH}][\mathrm{PIP_2}] - \mathrm{DF}k_{on}^{\mathrm{PH+PIP}}[\mathrm{PH.Arf}][\mathrm{PIP_2}] - \mathrm{DF}k_{on}^{\mathrm{PH+PIP}}[\mathrm{PH.Arf.ZA}][\mathrm{PIP_2}]$$

$$\frac{\mathrm{d[PH]}}{\mathrm{d}t} = k_{cat}^{\mathrm{PH.Arf.ZA}}[\mathrm{PH.Arf.ZA}] + k_{off}^{\mathrm{Arf+PH}}[\mathrm{PH.Arf}] + k_{off}^{\mathrm{Arf+PH}}[\mathrm{PH.Arf.ZA}] + k_{off}^{\mathrm{PH+PIP}}[\mathrm{PH_m}] - k_{on}^{\mathrm{Arf+PH}}[\mathrm{PH}][\mathrm{Arf}] - k_{on}^{\mathrm{Arf+PH}}[\mathrm{PH}][\mathrm{Arf.ZA}] - k_{on}^{\mathrm{PH+PIP}}[\mathrm{PH}][\mathrm{PIP_2}]$$

$$\frac{\mathrm{d[PH_m]}}{\mathrm{d}t} = k_{off}^{\mathrm{Arf+PH}}[\mathrm{PH_m.Arf.ZA}] + k_{off}^{\mathrm{Arf+PH}}[\mathrm{PH_m.Arf}] - k_{off}^{\mathrm{PH+PIP}}[\mathrm{PH_m}] + k_{cat}^{\mathrm{PH_m.Arf.ZA}}[\mathrm{PH_m.Arf.ZA}] + k_{on}^{\mathrm{PH+PIP}}[\mathrm{PH}][\mathrm{PIP_2}] - \mathrm{DF}k_{on}^{\mathrm{Arf+PH}}[\mathrm{PH_m}][\mathrm{Arf}] - \mathrm{DF}k_{on}^{\mathrm{Arf+PH}}[\mathrm{PH_m}][\mathrm{Arf.ZA}]$$

$$\frac{\mathrm{d[Arf]}}{\mathrm{d}t} = k_{off}^{\mathrm{Arf+ZA}}[\mathrm{Arf.ZA}] + k_{off}^{\mathrm{Arf+PH}}[\mathrm{PH.Arf}] + k_{off}^{\mathrm{Arf+PH}}[\mathrm{PH_m.Arf}] - k_{on}^{\mathrm{Arf+ZA}}[\mathrm{Arf}][\mathrm{ZA}] - k_{on}^{\mathrm{Arf+PH}}[\mathrm{PH}][\mathrm{Arf}] - \mathrm{DF}k_{on}^{\mathrm{Arf+PH}}[\mathrm{PH_m}][\mathrm{Arf}]$$

$$\frac{\mathrm{d[ZA]}}{\mathrm{d}t} = k_{cat}^{\mathrm{ZA}}[\mathrm{Arf.ZA}] + k_{off}^{\mathrm{Arf+ZA}}[\mathrm{PH_m.Arf.ZA}] + k_{off}^{\mathrm{Arf+ZA}}[\mathrm{PH.Arf.ZA}] + k_{off}^{\mathrm{Arf+ZA}}[\mathrm{Arf.ZA}] + k_{cat}^{\mathrm{PH.Arf.ZA}}[\mathrm{PH.Arf.ZA}] + k_{cat}^{\mathrm{PH_m.Arf.ZA}}[\mathrm{PH_m.Arf.ZA}] - k_{on}^{\mathrm{Arf+ZA}}[\mathrm{PH.Arf}][\mathrm{ZA}] - k_{on}^{\mathrm{Arf+ZA}}[\mathrm{Arf}][\mathrm{ZA}] - k_{on}^{\mathrm{Arf+ZA}}[\mathrm{ZA}][\mathrm{PH_m.Arf}]$$

$$\frac{\mathrm{d[Arf.ZA]}}{\mathrm{d}t} = -k_{cat}^{\mathrm{ZA}}[\mathrm{Arf.ZA}] - k_{off}^{\mathrm{Arf+ZA}}[\mathrm{Arf.ZA}] + k_{off}^{\mathrm{Arf+PH}}[\mathrm{PH_m.Arf.ZA}] + k_{off}^{\mathrm{Arf+PH}}[\mathrm{PH.Arf.ZA}] + k_{on}^{\mathrm{Arf+ZA}}[\mathrm{Arf}][\mathrm{ZA}] - k_{on}^{\mathrm{Arf+PH}}[\mathrm{PH}][\mathrm{Arf.ZA}] - \mathrm{DF}k_{on}^{\mathrm{Arf+PH}}[\mathrm{PH_m}][\mathrm{Arf.ZA}]$$

$$\frac{\mathrm{d[PH.Arf]}}{\mathrm{d}t} = k_{off}^{\mathrm{Arf+ZA}}[\mathrm{PH.Arf.ZA}] - k_{off}^{\mathrm{Arf+PH}}[\mathrm{PH.Arf}] + k_{off}^{\mathrm{PH+PIP}}[\mathrm{PH_m.Arf}] - k_{on}^{\mathrm{Arf+ZA}}[\mathrm{PH.Arf}][\mathrm{ZA}] + k_{on}^{\mathrm{Arf+PH}}[\mathrm{PH}][\mathrm{Arf}] - \mathrm{DF}k_{on}^{\mathrm{PH+PIP}}[\mathrm{PH.Arf}][\mathrm{PIP_2}]$$

$$\frac{\mathrm{d[PH.Arf.ZA]}}{\mathrm{d}t} = -k_{off}^{\mathrm{Arf+ZA}}[\mathrm{PH.Arf.ZA}] - k_{cat}^{\mathrm{PH.Arf.ZA}}[\mathrm{PH.Arf.ZA}] - k_{off}^{\mathrm{Arf+PH}}[\mathrm{PH.Arf.ZA}] + k_{off}^{\mathrm{PH+PIP}}[\mathrm{PH_m.Arf.ZA}] + k_{on}^{\mathrm{Arf+ZA}}[\mathrm{PH.Arf}][\mathrm{ZA}] + k_{on}^{\mathrm{Arf+PH}}[\mathrm{PH}][\mathrm{Arf.ZA}] - \mathrm{DF}k_{on}^{\mathrm{PH+PIP}}[\mathrm{PH.Arf.ZA}][\mathrm{PIP_2}]$$

$$\frac{\mathrm{d[PH_m.Arf]}}{\mathrm{d}t} = k_{off}^{\mathrm{Arf+ZA}}[\mathrm{PH_m.Arf.ZA}] - k_{off}^{\mathrm{Arf+PH}}[\mathrm{PH_m.Arf}] - k_{off}^{\mathrm{PH+PIP}}[\mathrm{PH_m.Arf}] - k_{on}^{\mathrm{Arf+ZA}}[\mathrm{ZA}][\mathrm{PH_m.Arf}] + \mathrm{DF}k_{on}^{\mathrm{Arf+PH}}[\mathrm{PH_m}][\mathrm{Arf}] + \mathrm{DF}k_{on}^{\mathrm{PH+PIP}}[\mathrm{PH.Arf}][\mathrm{PIP_2}]$$

$$\frac{\mathrm{d[PH_m.Arf.ZA]}}{\mathrm{d}t} = -k_{off}^{\mathrm{Arf+ZA}}[\mathrm{PH_m.Arf.ZA}] - k_{off}^{\mathrm{Arf+PH}}[\mathrm{PH_m.Arf.ZA}] - k_{off}^{\mathrm{PH+PIP}}[\mathrm{PH_m.Arf.ZA}] - k_{cat}^{\mathrm{PH_m.Arf.ZA}}[\mathrm{PH_m.Arf.ZA}] + k_{on}^{\mathrm{Arf+ZA}}[\mathrm{ZA}][\mathrm{PH_m.Arf}] + \mathrm{DF}k_{on}^{\mathrm{Arf+PH}}[\mathrm{PH_m}][\mathrm{Arf.ZA}] + \mathrm{DF}k_{on}^{\mathrm{PH+PIP}}[\mathrm{PH.Arf.ZA}][\mathrm{PIP_2}]$$

$$\frac{\mathrm{d[ArfGDP]}}{\mathrm{d}t} = k_{cat}^{\mathrm{ZA}}[\mathrm{Arf.ZA}] + k_{cat}^{\mathrm{PH.Arf.ZA}}[\mathrm{PH.Arf.ZA}] + k_{cat}^{\mathrm{PH_m.Arf.ZA}}[\mathrm{PH_m.Arf.ZA}]$$

### 2 PZA "tandem" Reaction ODEs

$$\begin{aligned}
\frac{\mathrm{d[Arf]}}{\mathrm{d}t} &= k_{off}^{\mathrm{Arf+ZA}}[\mathrm{Arf.ZA_m}] + k_{off}^{\mathrm{Arf+ZA}}[\mathrm{Arf.ZA}] + k_{off}^{\mathrm{Arf+PH}}[\mathrm{PH.Arf}] + k_{off}^{\mathrm{Arf+PH}}[\mathrm{PH_m.Arf}] \\
&\quad - k_{on}^{\mathrm{Arf+ZA}}[\mathrm{PZA}][\mathrm{Arf}] - k_{on}^{\mathrm{Arf+PH}}[\mathrm{PZA}][\mathrm{Arf}] - k_{on}^{\mathrm{Arf+ZA}}\mathrm{DF}[\mathrm{Arf}][\mathrm{PZA_m}] - \mathrm{DF}k_{on}^{\mathrm{Arf+PH}}[\mathrm{Arf}][\mathrm{PZA_m}] \\
\frac{\mathrm{d[PZA]}}{\mathrm{d}t} &= k_{cat}^{\mathrm{ZA}}[\mathrm{Arf.ZA}] + k_{off}^{\mathrm{Arf+ZA}}[\mathrm{Arf.ZA}] + k_{off}^{\mathrm{Arf+PH}}[\mathrm{PH.Arf}] + k_{cat}^{\mathrm{PH.Arf.ZA}}[\mathrm{PH.Arf.ZA}] \\
&\quad + k_{off}^{\mathrm{PH+PIP}}[\mathrm{PZA_m}] - k_{on}^{\mathrm{Arf+ZA}}[\mathrm{PZA}][\mathrm{Arf}] - k_{on}^{\mathrm{Arf+PH}}[\mathrm{PZA}][\mathrm{Arf}] - k_{on}^{\mathrm{PH+PIP}}[\mathrm{PZA}][\mathrm{PIP_2}] \\
\frac{\mathrm{d[PH.Arf]}}{\mathrm{d}t} &= k_{off}^{\mathrm{Arf+ZA}}[\mathrm{PH.Arf.ZA}] - k_{off}^{\mathrm{Arf+PH}}[\mathrm{PH.Arf}] + k_{off}^{\mathrm{PH+PIP}}[\mathrm{PH_m.Arf}] - x_{\mathrm{coop}}k_{on}^{\mathrm{Arf+ZA}}[\mathrm{PH.Arf}] \\
&\quad + k_{on}^{\mathrm{Arf+PH}}[\mathrm{PZA}][\mathrm{Arf}] - \mathrm{DF}k_{on}^{\mathrm{PH+PIP}}[\mathrm{PH.Arf}][\mathrm{PIP_2}] \\
\frac{\mathrm{d[PIP_2]}}{\mathrm{d}t} &= k_{off}^{\mathrm{PH+PIP}}[\mathrm{Arf.ZA_m}] + k_{off}^{\mathrm{PH+PIP}}[\mathrm{PH_m.Arf}] + k_{off}^{\mathrm{PH+PIP}}[\mathrm{PZA_m}] + k_{off}^{\mathrm{PH+PIP}}[\mathrm{PH_m.Arf.ZA}] \\
&\quad - k_{on}^{\mathrm{PH+PIP}}[\mathrm{PZA}][\mathrm{PIP_2}] - \mathrm{DF}k_{on}^{\mathrm{PH+PIP}}[\mathrm{PH.Arf}][\mathrm{PIP_2}] - \mathrm{DF}k_{on}^{\mathrm{PH+PIP}}[\mathrm{PH.Arf.ZA}][\mathrm{PIP_2}] \\
&\quad - \mathrm{DF}k_{on}^{\mathrm{PH+PIP}}[\mathrm{PIP_2}][\mathrm{Arf.ZA}] \\
\frac{\mathrm{d[PZA_m]}}{\mathrm{d}t} &= k_{cat}^{\mathrm{ZA}}[\mathrm{Arf.ZA_m}] + k_{off}^{\mathrm{Arf+ZA}}[\mathrm{Arf.ZA_m}] + k_{cat}^{\mathrm{PHm.Arf.ZA}}[\mathrm{PH_m.Arf.ZA}] + k_{off}^{\mathrm{Arf+PH}}[\mathrm{PH_m.Arf}] \\
&\quad - k_{off}^{\mathrm{PH+PIP}}[\mathrm{PZA_m}] + k_{on}^{\mathrm{PH+PIP}}[\mathrm{PZA}][\mathrm{PIP_2}] - k_{on}^{\mathrm{Arf+ZA}}\mathrm{DF}[\mathrm{Arf}][\mathrm{PZA_m}] - \mathrm{DF}k_{on}^{\mathrm{Arf+PH}}[\mathrm{Arf}][\mathrm{PZA_m}] \\
\frac{\mathrm{d[Arf.ZA]}}{\mathrm{d}t} &= -k_{cat}^{\mathrm{ZA}}[\mathrm{Arf.ZA}] - k_{off}^{\mathrm{Arf+ZA}}[\mathrm{Arf.ZA}] + k_{off}^{\mathrm{Arf+PH}}[\mathrm{PH.Arf.ZA}] + k_{off}^{\mathrm{PH+PIP}}[\mathrm{Arf.ZA_m}] \\
&\quad + k_{on}^{\mathrm{Arf+ZA}}[\mathrm{PZA}][\mathrm{Arf}] - x_{\mathrm{coop}}k_{on}^{\mathrm{Arf+PH}}[\mathrm{Arf.ZA}] - \mathrm{DF}k_{on}^{\mathrm{PH+PIP}}[\mathrm{PIP_2}][\mathrm{Arf.ZA}] \\
\frac{\mathrm{d[PH_m.Arf]}}{\mathrm{d}t} &= k_{off}^{\mathrm{Arf+ZA}}[\mathrm{PH_m.Arf.ZA}] - k_{off}^{\mathrm{Arf+PH}}[\mathrm{PH_m.Arf}] - k_{off}^{\mathrm{PH+PIP}}[\mathrm{PH_m.Arf}] - x_{\mathrm{coop}}k_{on}^{\mathrm{Arf+ZA}}[\mathrm{PH_m.Arf}] \\
&\quad + \mathrm{DF}k_{on}^{\mathrm{Arf+PH}}[\mathrm{Arf}][\mathrm{PZA_m}] + \mathrm{DF}k_{on}^{\mathrm{PH+PIP}}[\mathrm{PH.Arf}][\mathrm{PIP_2}] \\
\frac{\mathrm{d[PH.Arf.ZA]}}{\mathrm{d}t} &= -k_{off}^{\mathrm{Arf+ZA}}[\mathrm{PH.Arf.ZA}] - k_{off}^{\mathrm{Arf+PH}}[\mathrm{PH.Arf.ZA}] - k_{cat}^{\mathrm{PH.Arf.ZA}}[\mathrm{PH.Arf.ZA}] \\
&\quad + k_{off}^{\mathrm{PH+PIP}}[\mathrm{PH_m.Arf.ZA}] + x_{\mathrm{coop}}k_{on}^{\mathrm{Arf+ZA}}[\mathrm{PH.Arf}] + x_{\mathrm{coop}}k_{on}^{\mathrm{Arf+PH}}[\mathrm{Arf.ZA}] \\
&\quad - \mathrm{DF}k_{on}^{\mathrm{PH+PIP}}[\mathrm{PH.Arf.ZA}][\mathrm{PIP_2}] \\
\frac{\mathrm{d[PH_m.Arf.ZA]}}{\mathrm{d}t} &= -k_{off}^{\mathrm{Arf+ZA}}[\mathrm{PH_m.Arf.ZA}] - k_{cat}^{\mathrm{PHm.Arf.ZA}}[\mathrm{PH_m.Arf.ZA}] \\
&\quad - k_{off}^{\mathrm{Arf+PH}}[\mathrm{PH_m.Arf.ZA}] - k_{off}^{\mathrm{PH+PIP}}[\mathrm{PH_m.Arf.ZA}] \\
&\quad + x_{\mathrm{coop}}k_{on}^{\mathrm{Arf+ZA}}[\mathrm{PH_m.Arf}] + x_{\mathrm{coop}}k_{on}^{\mathrm{Arf+PH}}[\mathrm{Arf.ZA_m}] + \mathrm{DF}k_{on}^{\mathrm{PH+PIP}}[\mathrm{PH.Arf.ZA}][\mathrm{PIP_2}] \\
\frac{\mathrm{d[Arf.ZA_m]}}{\mathrm{d}t} &= -k_{cat}^{\mathrm{ZA}}[\mathrm{Arf.ZA_m}] - k_{off}^{\mathrm{Arf+ZA}}[\mathrm{Arf.ZA_m}] + k_{off}^{\mathrm{Arf+PH}}[\mathrm{PH_m.Arf.ZA}] - k_{off}^{\mathrm{PH+PIP}}[\mathrm{Arf.ZA_m}] \\
&\quad - x_{\mathrm{coop}}k_{on}^{\mathrm{Arf+PH}}[\mathrm{Arf.ZA_m}] + k_{on}^{\mathrm{Arf+ZA}}\mathrm{DF}[\mathrm{Arf}][\mathrm{PZA_m}] + \mathrm{DF}k_{on}^{\mathrm{PH+PIP}}[\mathrm{PIP_2}][\mathrm{Arf.ZA}] \\
\frac{\mathrm{d[ArfGDP]}}{\mathrm{d}t} &= k_{cat}^{\mathrm{ZA}}[\mathrm{Arf.ZA_m}] + k_{cat}^{\mathrm{ZA}}[\mathrm{Arf.ZA}] + k_{cat}^{\mathrm{PHm.Arf.ZA}}[\mathrm{PH_m.Arf.ZA}] + k_{cat}^{\mathrm{PH.Arf.ZA}}[\mathrm{PH.Arf.ZA}]
\end{aligned}$$

**Supplementary References**


1.Bezanson, J., Edelman, A., Karpinski, S. & Shah, V. B. Julia: A Fresh Approach to Numerical Computing. SIAM Rev. 59, 65–98 (2017).

2.Rackauckas, C. & Nie, Q. DifferentialEquations.jl – A Performant and Feature-Rich Ecosystem for Solving Differential Equations in Julia. Journal of Open Research Software 5, (2017).

3.Art et al. Wildart/Evolutionary.Jl: V0.11.1. (Zenodo, 2022). doi:10.5281/zenodo.5851574.

4.Loman, T. E. et al. Catalyst: Fast and flexible modeling of reaction networks. PLOS Computational Biology 19, e1011530 (2023).

5.Wu, Y., Vendome, J., Shapiro, L., Ben-Shaul, A. & Honig, B. Transforming binding affinities from three dimensions to two with application to cadherin clustering. Nature 475, 510–513 (2011).

6.Jian, X. et al. Molecular Basis for Cooperative Binding of Anionic Phospholipids to the PH Domain of the Arf GAP ASAP1. Structure 23, 1977–1988 (2015).

7.Luo, R. et al. Kinetic analysis of GTP hydrolysis catalysed by the Arf1-GTP–ASAP1 complex. Biochemical Journal 402, 439–447 (2007).
